# Identification of G Protein α_i_ Signaling Partners by Proximity Labeling Reveals a Network of Interactions that Includes PDZ-RhoGEF

**DOI:** 10.1101/2021.07.15.452545

**Authors:** Naincy R. Chandan, Saji Abraham, Shuvasree SenGupta, Carole A. Parent, Alan V. Smrcka

**Affiliations:** Department of Pharmacology, University of Michigan, Ann Arbor, MI 48109, USA; Department of Cell and Developmental Biology, University of Michigan, Ann Arbor, MI 48109, USA; Rogel Cancer Center Michigan Medicine, University of Michigan, Ann Arbor, MI 48109, USA; Life Sciences Institute, University of Michigan, Ann Arbor, MI, 48109, USA

**Author notes:** Corresponding author: Alan V. Smrcka, **Email:**.

**Keywords:** GPCR signaling, Gα_i_ proteins, Proximity labeling, Mass spectrometry, Interactome, PDZ-RhoGEF, Cell-migration

## Abstract

G protein-coupled receptors (GPCRs) that couple to the Gi family of G proteins are key regulators of cell and tissue physiology. Our recent work has discovered novel roles for Gα_i_ in migration of neutrophils and fibrosarcoma cells downstream of activated chemoattractant receptors, but the molecular target(s) of Gα_i_ in these processes remain to be identified. We adopted an intact cell proximity-based labeling approach using BioID2 coupled to tandem mass tag (TMT)-based quantitative proteomics to identify proteins that selectively interact with the GTP-bound form of Gα_i1_. Multiple targets were identified and validated for selective biotinylation by active BioID2-Gα_i1_(Q204L), suggesting a previously unappreciated network of interactions for activated Gα_i_ proteins in intact cells. Extensive characterization of one candidate protein, PDZ-RhoGEF (PRG), revealed that active-Gα_i1_ strongly activates PRG. Strikingly, large differences in the ability of Gα_i1_, Gα_i2_, and Gα_i3_ isoforms to activate PRG were observed despite over 85% sequence identity. We also demonstrate the functional relevance of the interaction between active Gα_i_ and PRG *ex vivo* in primary human neutrophils. Identification and characterization of new targets regulated by Gα_i_ both individually and in networks provide insights that will aid not only in investigation of diverse functional roles of Gi-coupled GPCRs in biology but also in the development of novel therapeutic approaches.

**Summary:** Proximity-based labeling approach was used to identify signaling networks and signaling mechanisms downstream of Gi-coupled receptors.

## Introduction

G protein-coupled receptors (GPCRs) are a major class of cell surface receptors that regulate a variety of physiological and pathophysiological processes in response to a wide variety of ligands, including neurotransmitters, chemoattractants, hormones, opioids, and drugs. Activated GPCRs bind to heterotrimeric G proteins consisting of Gα subunits and Gβγ constitutive heterodimers and catalyze the exchange of GTP for GDP on the Gα subunits. Subsequent conformational changes in the Gα subunit cause it to dissociate from the receptor and Gβγ subunits. Both Gα and Gβγ subunits then transduce signals from receptors to a range of downstream effector proteins, including second messenger generating enzymes and ion channels [1–4].

Gα_i/o_ coupled GPCRs comprise nearly one hundred receptors for a wide variety of ligands including opioids, cannabinoids, prostaglandins, histamine, somatostatins, chemokines, and neurotransmitters such as acetylcholine, adrenaline, serotonin, and dopamine [5]. The α-subunits that define the basic properties of heterotrimeric G proteins are divided into four families, Gα_s_, Gα_i/o_, Gα_q/11_, and Gα_12/13_ [1]. The Gα_i/o_ family is an abundant and ubiquitous class of G protein subunits consisting of various isoforms including Gα_i1_, Gα_i2_, Gα_i3_, and Gα_o_ [5]. Gα_i/o_ subunit activity is classically associated with adenylate cyclase (AC) inhibition [6].

Gα_i/o_-coupled chemokine and chemoattractant GPCRs regulate directional cell migration and adhesion, which is involved in tissue formation, wound healing, immune responses, and cancer cell invasion and metastasis [7–9]. It is well accepted that Gβγ subunits released from Gi heterotrimers are central mediators of chemokine driven chemotaxis, while Gα_i_ has been proposed to function passively through the GDP-GTP exchange-dependent cycling of free and bound Gβγ subunits [10]. Identification of signaling mechanisms specifically downstream of Gα_i_ subunits has been hampered by the fact that perturbations that inhibit Gα_i_ signaling also inactivate Gβγ signaling. For example, modification of Gα_i_ by pertussis toxin (PTX) blocks interactions between the Gα_i_-βγ heterotrimer and GPCRs, thereby inhibiting both Gα and Gβγ signaling [11]. Similarly, knockout (KO) of specific G protein α subunits, either in mice or with specific short inhibitory RNAs in cell culture, prevents signaling by both Gα and its associated Gβγ subunits [12]. Using a small-molecule Gβγ activator developed in our laboratory, we identified a role for active Gα_i_ (Gα_i_-GTP) in regulating neutrophil and HT1080 fibrosarcoma cell migration [13, 14]. In these studies, we showed that Gβγ promotes cell adhesion and Gα_i_-GTP promotes de-adhesion, processes that must be coordinated for cells to move [14]. Gα_i_-GTP regulation of adhesion was independent of cAMP signaling [13]; however, a direct effector regulated by Gα_i_ was not identified.

Methods previously used to identify new effectors of Gα_i_ beyond AC include the yeast-two hybrid system and immunoprecipitation (IP) followed by mass spectrometry (MS) [15–17]. While these methods have successfully identified interacting proteins, they have limitations. IP-MS methods recover only strong interaction partners that survive cell lysis and repeated detergent washes. GPCR-dependent signal transduction processes often involve transient protein-protein interactions that are lost after cell disruption. The yeast two-hybrid systems lack appropriate cellular context, and only fragments of proteins are used to identify binding interactions. It is clear that cell context is critical for optimizing interactions between signal transduction components through compartmentalization and interactions with membrane surfaces. Thus, it is likely that multiple G protein interactions may have been missed by these traditional approaches.

To circumvent these challenges, we adopted a proximity labeling approach using BioID2, a promiscuous biotin ligase enzyme, coupled to affinity purification and MS [18, 19]. Our goal was to capture Gα_i_ subunit interactions with novel potential signal transduction partners and complexes in the context of intact cells of interest. Using this approach, we identified multiple known Gα_i_ binding partners, including Gβγ subunits and AC. Multiple classes of proteins involved in diverse cellular processes, including cell migration and amino acid transport, were identified as potential interaction partners of active Gα_i1_. One of the proteins identified was the RH family RhoGEF, PDZ-RhoGEF (PRG), also known as ARHGEF11. The role of PRG in regulation of cell migration downstream of Gi-coupled chemoattractant receptors is well characterized, but it is thought to be mediated primarily by additional coupling of these receptors to Gα_12/13_ subunits [20, 21]. We also demonstrate that PRG is an effector of active Gα_i1_ and Gα_i3_ but is poorly activated by Gα_i2_, a highly homologous (~85% identical) Gα_i_ family member. We demonstrate that PRG can be activated downstream of Gi-coupled receptors and show Gα_i_’s involvement in regulation of PRG in human neutrophils. PRG is relatively ubiquitously expressed [22]; thus, its identification as a new Gα_i_-GTP target has implications for regulation of Rho in various tissues and cell types by Gi-coupled GPCRs.

## Results

### Rational Design of BioID2 Fused Gα_i1_

To identify proteins that selectively interact with the active form of Gα_i1_ using proximity labeling, we fused the promiscuous biotin ligase, BioID2 [19], to Gα_i1_ (BioID2-Gα_i1_) and constitutively active Gα_i1_-Q204L (BioID2-Gα_i1_-QL). We inserted BioID2 as an internal tag in the αb-αc loop of Gα_i1_, which has been shown to tolerate GFP insertion allowing the N and C termini to interact with membranes and receptors [23] (Fig. 1A). In the absence of receptor-dependent activation in cells, Gα_i1_ is primarily GDP-bound and inactive (referred to as Gα_i1_), whereas a Q204L mutation in Gα_i1_ renders it GTPase-deficient, and therefore, GTP-bound and constitutively active (referred to as Gα_i1_-QL) [24]. Comparison of active versus inactive Gα_i1_ allowed us to search for targets that interact selectively with the activated conformation of Gα_i1_. As a control for general promiscuous labeling of proteins because of abundance or simply due to co-residence at the plasma membrane (PM), a PM-targeted BioID2 fused to the C-terminal PM targeting motif of KRas (BioID2-CaaX) was used. Thus, three experimental groups, BioID2-Gα_i1_, BioID2-Gα_i1_-QL, and BioID2-CaaX, were used to screen for potential targets that selectively interact with Gα_i1_-GTP (Fig. 1A left panel).

**Figure 1.**
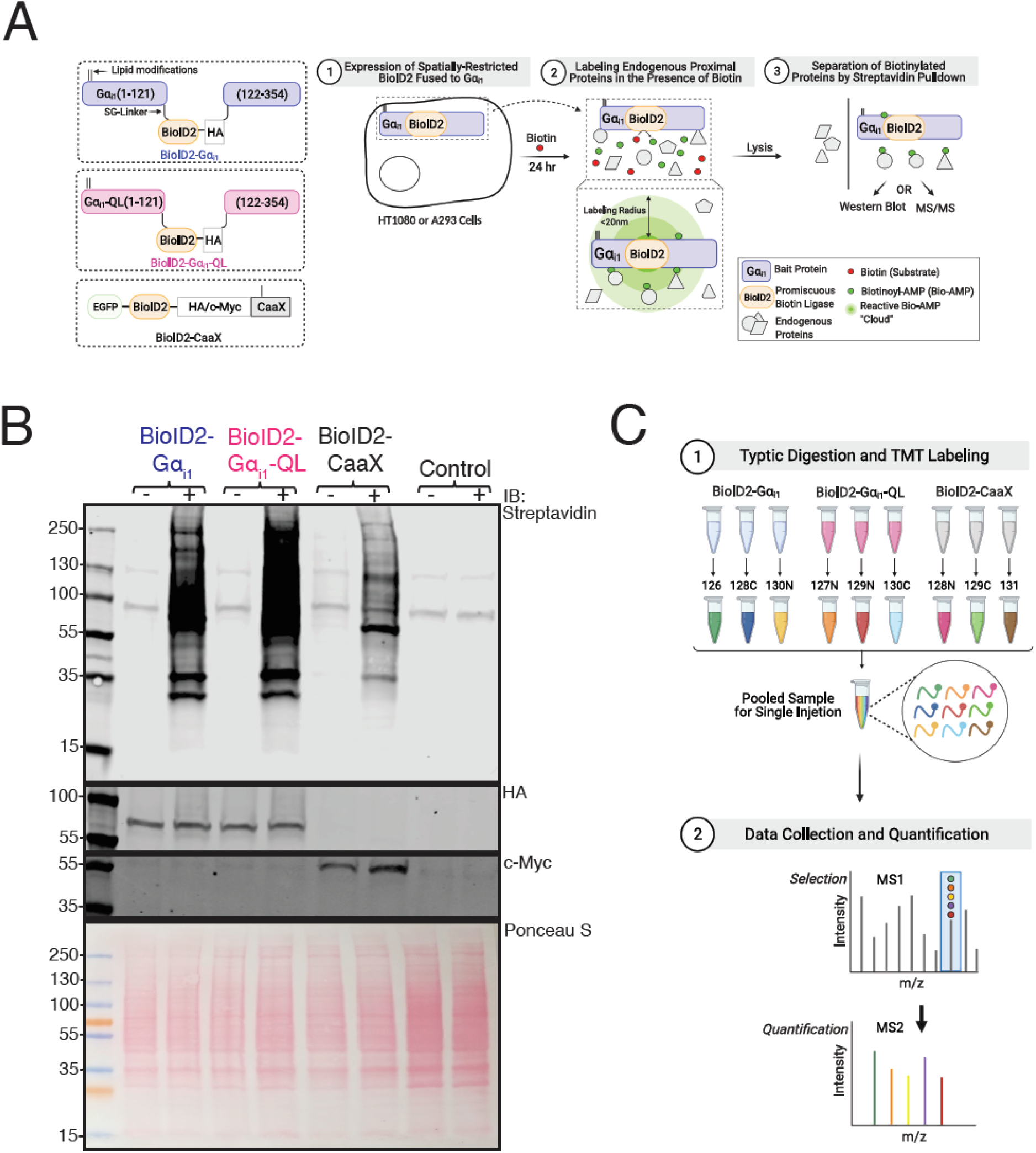
Principle and Experimental Workflow of Proximity Labeling of the Gα_i1_ Interactome. **(A)** Left Panel: Schematic of BioID2 fusion constructs. BioID2 was inserted between residues A121-E122 in the αb-αc loop (the first loop of the helical domain) of human Gα_i1_, flanked by SG-linkers. Palmitoylation and myristylation sites on the Gα_i1_ subunit and the farnesylation sites on CaaX moiety are labeled as lipid modifications. Right Panel: Schematic of principle and experimental workflow of proximity-based labeling using BioID2. HT1080 cells were used for mass spectrometry experiments and A293 cells were used for pull-down western blot. Cells were transfected with the indicated constructs and labeled for 24 hr in the presence of biotin. Gα_i1_ fused BioID2 biotinylates proteins in proximity (< 20 nm) in an unbiased manner to identify candidate interacting proteins of Gα_i1_. **(B)** Transfected BioID2 fused constructs biotinylate multiple proteins in cells. A293 cells were transfected with indicated constructs and labeled in the presence of biotin for 24 hr. Top panel: Biotinylated proteins present in the whole-cell lysates after 24 hr of labeling were detected on a streptavidin western blot. The two bands at 130 and ~90 kDa correspond to endogenously biotinylated proteins in control lanes. Middle panels: Expression of the BioID2-Gα_i1_ and QL was tested with Gα_i1/2_ antisera and BioID2-CaaX was tested using an anti-c-Myc antibody on western blots. Bottom Panel: Ponceau S-stained blot showing total protein loading. Western blots represent one of three independent experiments that yielded similar results. **(C)** Schematic of sample processing and mass spectrometry analysis. Samples pulled down using streptavidin beads were digested with trypsin and labelled with a TMT tag. Triplicate samples of BioID2-Gα_i1_ and BioID2-Gα_i1_-QL and BioID2-CaaX were pooled and resolved by LC-MS and the data was analyzed using proteome discover.

### BioID2 Fused Gα_i1_ Localizes Predominantly to the PM and Gα_i1_-QL Inhibits cAMP Accumulation

To characterize the functionality of BioID2 fused Gα_i1_ proteins, we examined their localization in A293 cells. All three proteins, BioID2-Gα_i1_, BioID2-Gα_i1_-QL, and BioID2-CaaX, localized predominantly to the PM (Fig. S1A). Next, we evaluated the ability of Gα_i1_-QL to inhibit AC by measuring inhibition of forskolin (Fsk)-stimulated cyclic-AMP (cAMP) production using a cAMP biosensor (cAMP-Glo™). As expected, both untagged and BioID2 tagged Gα_i1_-QL significantly reduced the rate and extent of cAMP generated upon Fsk addition (Fig. S1B). Conversely, the Gα_i1_-GDP counterparts did not affect Fsk stimulated cAMP accumulation. Western blot confirmed that expression of these proteins was equal (Fig. S1C). These experiments established that Gα_i1_-QL fused with BioID2 can localize to the PM, inhibit AC, and thus behave similarly to the untagged counterpart.

### BioID2 Fused with Gα_i1_ Biotinylates Endogenous Proteins

We tested the biotin labeling efficiency of BioID2 fused Gα_i1_ proteins in A293 cells. Cells were transiently transfected with the indicated cDNA clones and incubated with biotin for 24 hr (Fig. 1A right panel). Following whole-cell lysis, proteins were resolved by SDS-gel electrophoresis and probed with streptavidin-IRDye 800 on a western blot. Multiple proteins were biotinylated in BioID2-Gα_i1_, BioID2-Gα_i1_-QL, and BioID2-CaaX samples, and biotinylation was dependent on BioID2 and biotin (Fig. 1B). There were differences in the total biotinylation pattern amongst the three experimental groups, suggesting that the three fusion proteins label endogenous proteins differentially.

### Proximity Labeling-Coupled MS in HT1080 Fibrosarcoma Cells

One goal of these studies was to identify proteins regulated by Gα_i_ that could be involved in cell migration downstream of chemokine/chemoattractant receptors. HT1080 fibrosarcoma cells express FPR1 receptors, adhere and migrate on fibronectin-coated surfaces, and are comparatively easy to grow and transfect relative to neutrophil-like cells. Roles for Gα_i1_ and Gβγ in cell adhesion and migration have been previously established in these cells [14, 25]. For these reasons, we chose HT1080 cells for the proximity labeling experiments to increase the probability of identifying effectors of Gα_i1_ relevant to cell migration.

Three sets of transfections with BioID2-Gα_i1_, BioID2-Gα_i1_-QL, and BioID2-CaaX, were independently performed in HT1080 cells, and BioID2 fused Gα_i1_ subunits were expressed at levels similar to endogenous Gα_i_ (Fig. S1D). To perform quantitative comparison of biotinylated proteins after purification with streptavidin beads, each sample was labeled with a unique isobaric tandem mass tag (TMT). This allowed triplicate samples for each group to be pooled and analyzed in a single MS run so that relative abundance of thousands of proteins could be compared (Fig. 1C).

### MS Data Analysis Identifies Multiple Candidate Interacting Proteins

We detected several proteins known to interact with Gα_i_ including, Gβ and γ subunit isoforms, which were selectively enriched in the BioID2-Gα_i1_ samples relative to BioID2-Gα_i1_-QL, as expected (Fig. 2A). Several isoforms of AC were detected, but there was no statistically significant difference observed between the BioID2-Gα_i1_-QL and BioID2-Gα_i1_ samples. Ric8A was also equally labeled by BioID2-Gα_i1_-QL and BioID2-Gα_i1_. Gα_i_-GTP effectors, GPRIN1 and RASA3 [16, 26], were highly enriched in BioID2-Gα_i1_-QL samples relative to BioID2-Gα_i1_ samples (Fig. 2A). A number of receptors were also identified, but most were not significantly enriched in either the BioID2-Gα_i1_-QL or BioID2-Gα_i1_ samples (Table S1).

**Figure 2.**
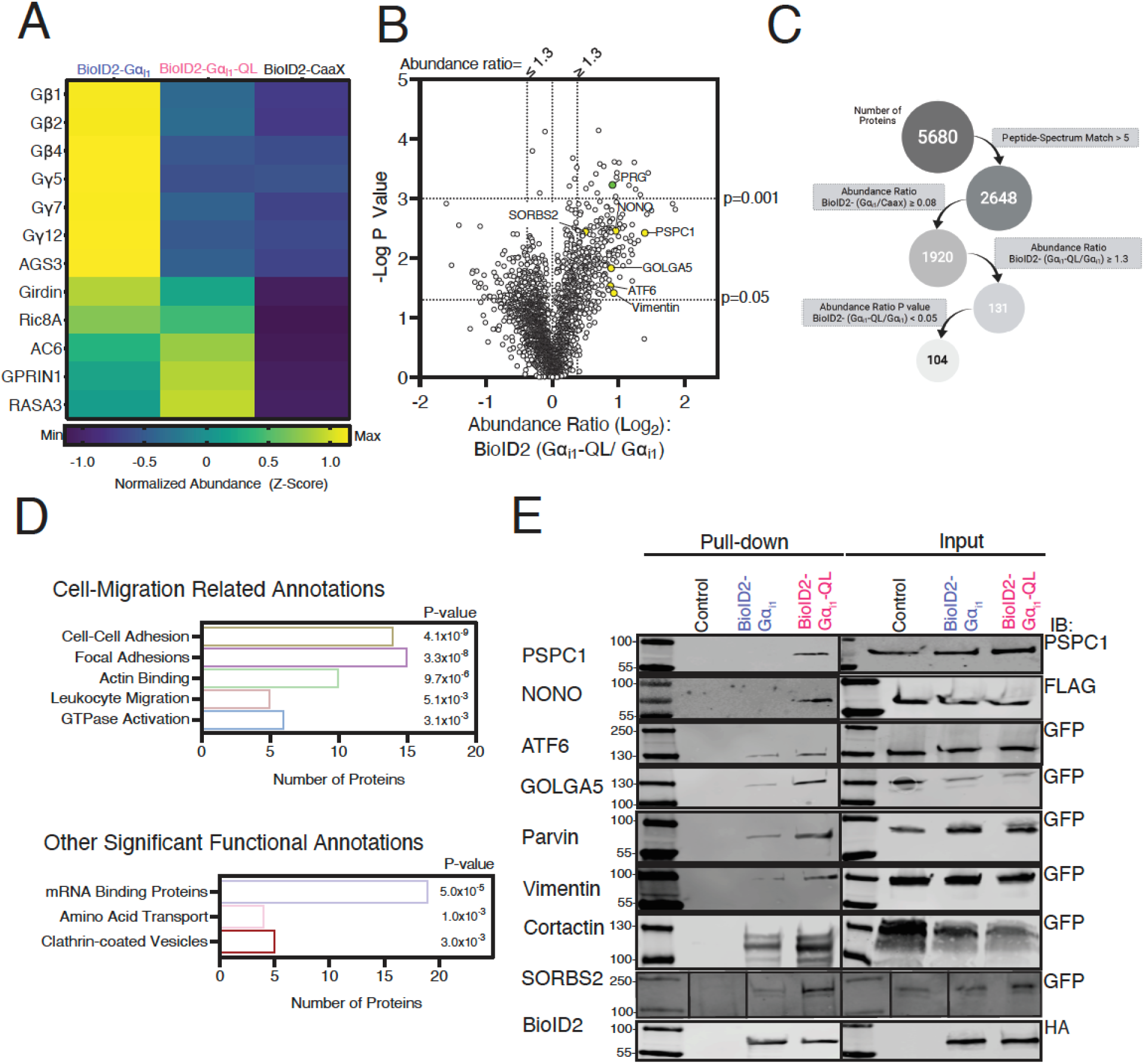
Proximity Labeling Proteomics Results. **(A)** Heat map showing the relative changes in abundance of known binding partners of Gα_i_ that were identified in the mass spectrometry analysis. **(B)** Volcano plot of all high confidence proteins identified where the BioID2-Gα_i1_/BioID2-CaaX ratio was greater than 0.8. PRG is highlighted in green, and the candidate proteins investigated in 2E are labeled in yellow. **(C)** Schematic showing filtering criteria for selection of proteins enriched in BioID2-Gα_i1_-QL samples relative to BioID2-Gα_i1_. **(D)** Representative classes from Go analysis of proteins from B that met the final criteria in C. P values were generated with the DAVID software. **(E)** Validation of candidate proteins for their proximity and enrichment with BioID2-Gα_i1_-QL. cDNA clones encoding indicated epitope or GFP tagged proteins were co-transfected with BioID2-Gα_i1_ or BioID2-Gα_i1_-QL, labeled with biotin for 24 hr, followed by streptavidin pull-down and western blotting. Western blots are representative of experiments performed twice yielding qualitatively comparable data.

Overall, ~5000 proteins were isolated and identified. We selected proteins with a minimum of 5 peptides assigned to each protein to ensure robustness of the data. We also filtered the data to only include proteins where the ratio of normalized abundance for the BioID2-Gα_i1_ and BioID2-CaaX is roughly equivalent or greater (BioID2-Gα_i1_/ BioID2-CaaX > 0.8). The rationale behind this was that since BioID2-CaaX labels proteins at the PM based on proximity within the compartment, proteins labeled equally by BioID2-CaaX and BioID2-Gα_i1_ are likely colocalized with BioID2-Gα_i1_ at PM. Proteins labeled to a greater extent by BioID2-Gα_i1_ than BioID2-CaaX could be PM resident proteins that selectively interact with Gα_i1_ in its inactive GDP-bound state but could also be proteins labeled by BioID2-Gα_i1_ in other compartments or cytosolic proteins that interact with Gα_i1_ at the PM. The data are shown as a volcano plot in Fig. 2B.

To identify proteins that selectively interact with Gα_i1_-QL, we further filtered the data and included only those proteins with BioID2-Gα_i1_-QL/BioID2-Gα_i1_ normalized abundance ratio ≥ 1.3 and a P-value < 0.05 (Fig. 2C). This resulted in a list of 104 candidate proteins (Fig. 2C, S2, Table S2). These 104 Gα_i1_-QL enriched proteins were analyzed using DAVID gene ontology software to identify classes of proteins involved in different cellular processes [27]. Several enriched targets regulate various aspects of cell migration (Fig. 2D top panel, Table S3). These data suggest that active Gα_i_ may regulate cell migration through a protein interaction network rather than just a single target. Other classes of proteins identified with high confidence were mRNA binding proteins, amino acid transporters, and proteins involved in clathrin-mediated endocytosis (Fig. 2D bottom panel, Table S3).

Most of these proteins have not been previously identified as targets of Gα_i_. To further validate selective enrichment of potential Gα_i_ binding proteins with BioID2-Gα_i1_-QL, we selected a set of enriched proteins based on the availability of epitope or fluorescent protein (FP)-tagged cDNA clones to test in a proximity labeling-coupled biotinylation western blot assay. Proteins tested in this assay were: PSPC1 (Paraspeckle Component 1), p54 (Nuclear RNA-Binding Protein, 54-KD or NONO), ATF6 (Activating Transcription Factor 6), SORBS (Sorbin and SH3 domain-containing protein 2 or ArgBP2), GOLGA5 (Golgin A5) and Vimentin (Fig. 2B, highlighted in yellow). We also included two proteins of interest that did not quite reach statistical significance, Parvin (BioID2-Gα_i1_-QL/BioID2-Gα_i1_=1.33, p=0.08) and Cortactin (BioID2-Gα_i1_-QL/BioID2-Gα_i1_=1.5, p=0.053).

Each protein-coding cDNA was individually co-transfected with BioID2-Gα_i1_-QL, BioID2-Gα_i1_, or control plasmid in A293 cells (Fig. 2E). Cells were treated with biotin, followed by cell lysis, streptavidin pull-down, and western blotting. Of the 12 cDNA clones tested, 8 showed enriched labeling by BioID2-Gα_i1_-QL relative to BioID2-Gα_i1_ (Fig. 2E), indicating that there is indeed preferential interaction between the active form of Gα_i1_ and the candidate proteins. Co-transfection of these cDNAs with BioID2 tagged Gα_i1_ cDNAs did not increase expression of the proteins, indicating that increased biotin labeling is not due to an increase in expression. These data support the idea that many of the other proteins amongst the 104 proteins enriched in the BioID2-Gα_i1_-QL samples are indeed in selective proximity to the active form of Gα_i1_.

Overall, the data suggest that Gα_i_ regulates multiple classes of cellular processes via mechanisms that involve coordinated network interactions with a variety of protein targets.

### PRG Selectively Interacts with Active Gα_i1_

One protein of interest relevant to cell migration and significantly enriched in BioID2-Gα_i1_-QL relative to BioID2-Gα_i1_ samples was PDZ-RhoGEF (PRG, or ARHGEF11), a Rho guanine nucleotide exchange factor (Fig. 2B, highlighted in green and 3A). PRG biotin labeling by BioID2-Gα_i1_ and BioID2-CaaX was roughly equivalent based on the MS quantification, suggesting labeling by inactive BioID2-Gα_i1_ was primarily due to membrane proximity (Fig. 3A). We decided to pursue PRG for several reasons; PRG was strongly enriched in the BioID2-Gα_i1_-QL samples with a P-value <0.001, and PRG has an established role in regulation of neutrophil migration downstream of Gi-coupled chemoattractant receptors [20, 21]. PRG localizes to the back of migrating neutrophils and activates Rho and myosin-dependent tail retraction during migration in response to chemoattractants [21]. To ensure that the high PRG abundance in BioID2-Gα_i1_-QL samples was not simply the result of increased PRG expression, we compared the expression of endogenous PRG in HT1080 cells transfected with BioID2-Gα_i1_, BioID2-Gα_i1_-QL, or BioID2-CaaX by western blotting. Endogenous PRG expression was equal in all three conditions (Fig. 3B).

**Figure 3.**
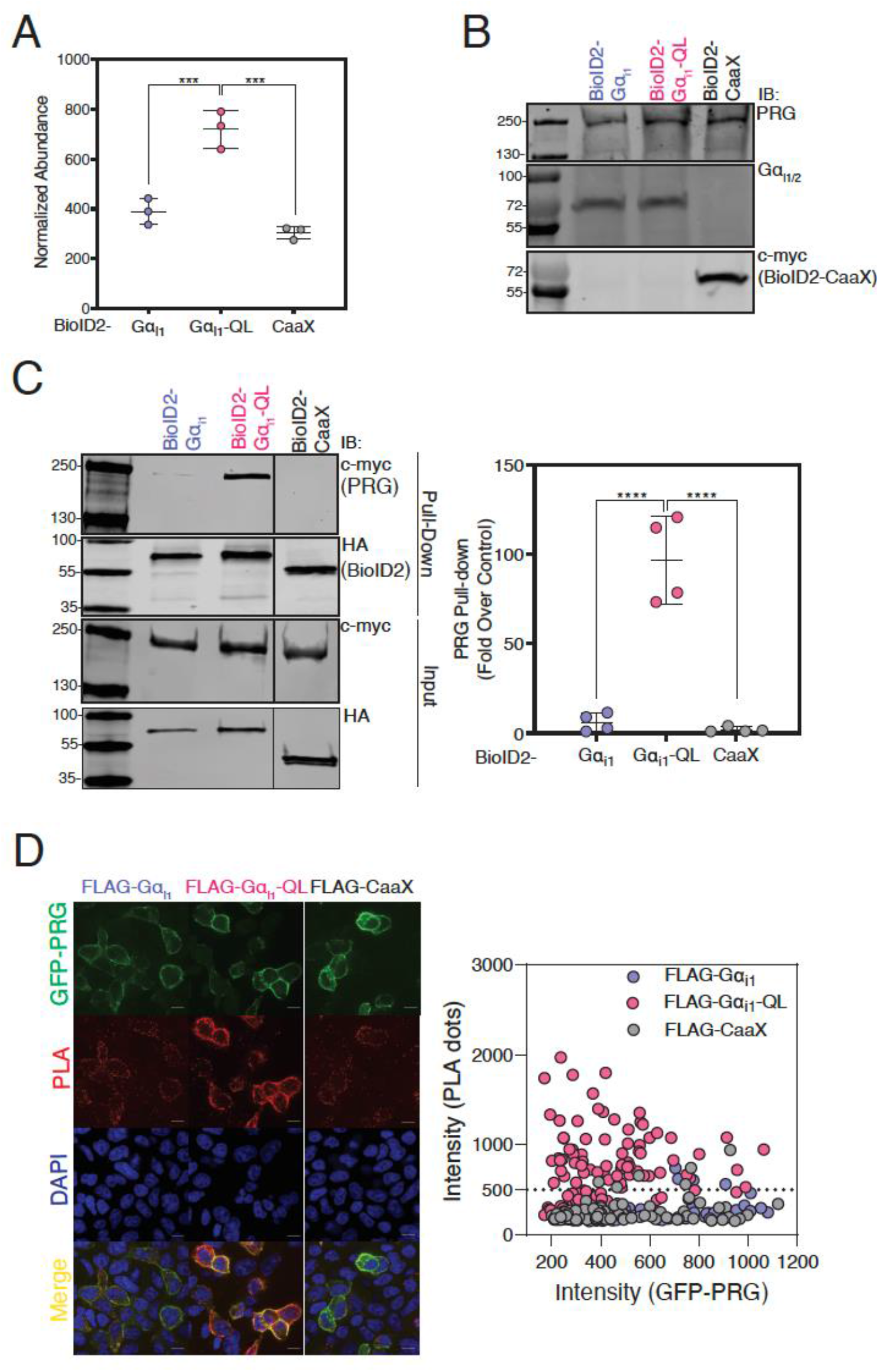
BioID2-Gα_i1_-QL Interacts with PRG in Cells. **(A)** Increased PRG abundance in BioID2-Gα_i1_-QL. Normalized abundance of PRG was quantified by MS. **(B)** Western blot showing equal endogenous PRG expression in BioID2-Gα_i1_, BioID2-Gα_i1_-QL or BioID2-CaaX expressing HT1080 cells. HT1080 cells were transfected with indicated constructs and WCLs were resolved on a western blot. **(C)** Enrichment of PRG biotinylation by BioID2-Gα_i1_-QL confirmed by biotin labeling, pull-down and western blotting. A293 cells were transfected with indicated cDNA constructs, labeled in the presence of biotin for 24 hr, and subsequently pulled down using streptavidin. Left panel: Representative western blots of PRG isolated using streptavidin pull-down and expression of BioID2-Gα_i1_, Gα_i1_-QL and BioID2-CaaX. Right Panel: Quantitation of three independent experiments normalized to total PRG (right panel). The data represent the mean ± SD of three separate experiments. (n=3, one-way ANOVA. ***P < 0.001, ****<0.0001). **(D)** GFP-PRG and FLAG-Gα_i1_-QL interact in a proximity ligation assay (PLA). Following transfection of APEX2-FLAG fused Gα_i1_-WT, Gα_i1_-QL, or CaaX with GFP-PRG constructs for 48 hr, cells were fixed, permeabilized, and subsequent PLA assay steps were followed as detailed in methods. Left panels: Three random confocal fields were imaged, and representative images are shown for GFP-PRG (green), PLA reaction (red), merge (orange) and DAPI (blue). Scale bar, 10μm. Right panel: The intensity of the PLA signal was quantitated (y-axis) and was plotted against GFP-PRG expression (x-axis). Control experiments omitting any one of the protein partners or primary antibodies resulted in no observable PLA signal (data not shown). For each experiment, ~100 cells per condition were analyzed, and the data is representative of one of three independent experiments that yielded similar results.

To independently validate PRG identification by MS, we co-expressed PRG with BioID2-Gα_i1_, BioID2-Gα_i1_-QL, or BioID2-CaaX in A293 cells, labeled with biotin, followed by streptavidin affinity pull-down and western blotting for PRG. PRG was highly enriched in the BioID2-Gα_i1_-QL samples compared to BioID2-Gα_i1_ and BioID2-CaaX (Fig. 3C).

Next, we used a proximity ligation assay (PLA) to test for interactions in A293 cells. A robust PLA signal was detected when GFP-PRG was co-transfected with APEX-FLAG-Gα_i1_-QL. The low PLA signal observed between GFP-PRG and controls APEX-FLAG-Gα_i1_-WT, or APEX-CaaX is likely due to colocalization at the PM, resulting in background bystander proximity labeling (Fig.3D). These data further support selective interactions between Gα_i1_-GTP and PRG (Fig. 3D).

### Gα_i1_ Activates RhoGEF Activity of PRG

These findings prompted us to investigate whether Gα_i1_ could activate PRG. An SRE luciferase (SRE-Luc) reporter, responsive to Rho activation [28], was used to study PRG activation in transfected A293 cells. We reconstituted the signaling pathway components by co-transfecting wild-type Gα_i1_ (Gα_i1_-WT) or constitutively active Gα_i1_ (Gα_i1_-QL) with PRG and the SRE-Luc reporter. PRG or Gα_i1_-QL, when transfected independently, did not stimulate SRE-Luc, but when transfected together, strong synergistic activation of SRE-Luc was observed (Fig. 4A left panel), indicating that Gα_i1_-QL activates PRG to stimulate Rho activation. Activation of PRG was dependent on the Gα_i1_ activation state since Gα_i1_-WT did not significantly increase PRG activity (Fig. 4A left panel). Similar results were seen with an SRF-Luc reporter (data not shown). PRG activation by Gα_i1_-QL was concentration-dependent (Fig. 4A middle panel). Gα_i1_-WT and Gα_i1_-QL were expressed at comparable levels, and PRG expression was similar in all three conditions assessed by western blotting (Fig. 4A right panel).

**Figure 4.**
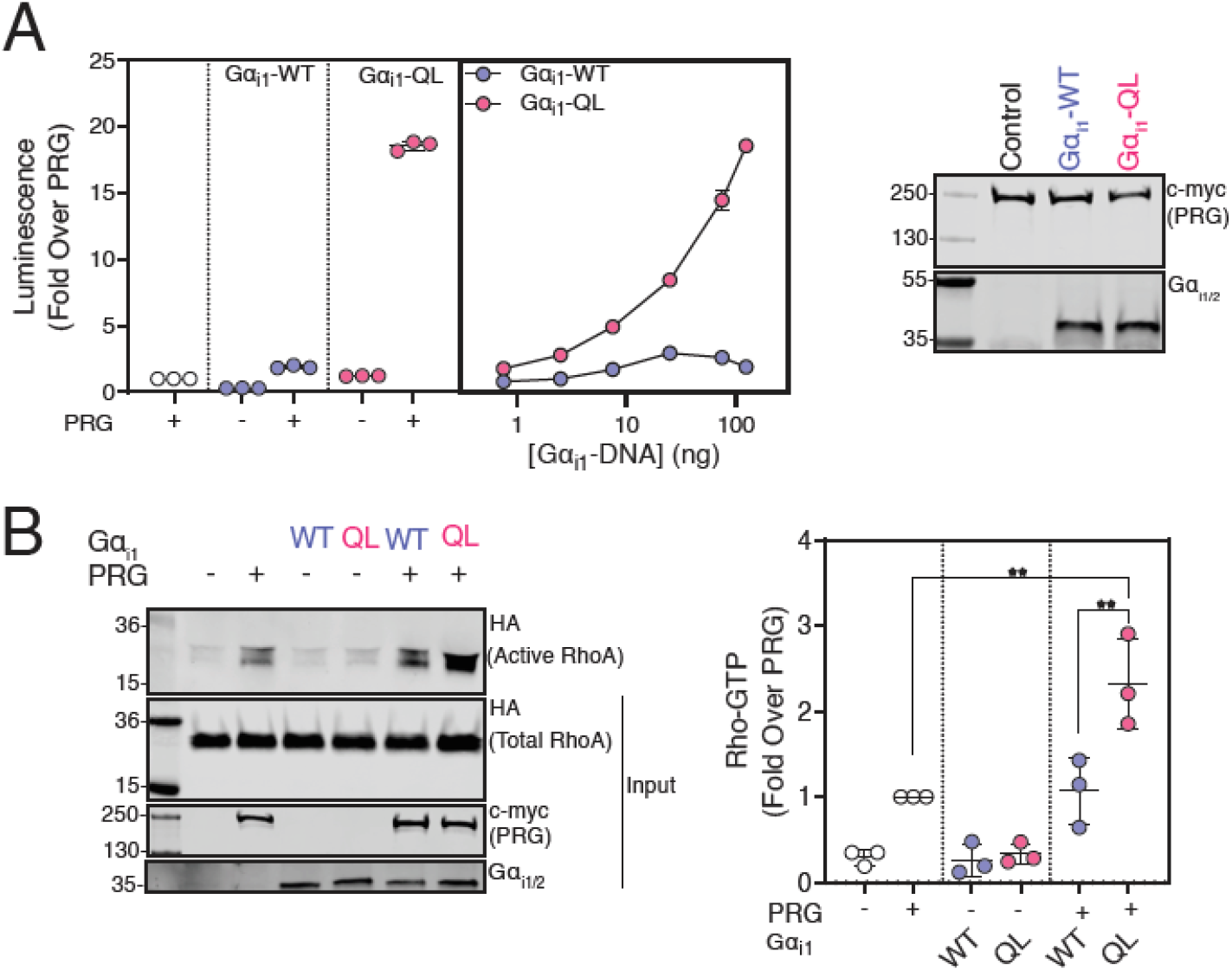
Gα_i1_-GTP Activates PRG Activity. **(A)** Gα_i1_-QL activates PRG in SRE-Luc reporter assay and the activation is dependent on the Gα_i1_ concentration. A293 cells were transfected with cDNAs encoding SRE-Luciferase, PRG, and either Gα_i1_ or Gα_i1_-QL for 20 hr. Left and middle Panels: Cells were serum-starved for 4 hr, and luminescence was measured 10 min after addition of One-Glo™ reagent. Data are representative of three independent experiments. Right panel: Representative western blots showing relative expression of various cDNA constructs in A293 cells. Data are representative of three independent experiments that yielded similar results. **(B)** Gα_i1_-QL increases active RhoA pull-down in a Rhotekin binding assay. A293 cells were transfected with indicated cDNA constructs for 24 hr, lysed, and incubated with GST-Rhotekin beads for 1 hr. Left panel: Representative western blots showing bound RhoA-GTP and relative expression of transfected constructs. Right panel: Quantitative comparison of three independent experiments, normalized to total RhoA. The data represent the mean ± SD of three separate experiments. (n=3, one-way ANOVA. **P < 0.01).

To validate activation of PRG by active Gα_i1_, we measured active RhoA (RhoA-GTP) in response to Gα_i1_-WT or Gα_i1_-QL co-expression with PRG in A293 cells with a Rhotekin pull-down assay. In this assay, RhoA binds to Rhotekin-Glutathione S-transferase (GST) in an activation-dependent manner, which can then be precipitated and immunoblotted. Expression of exogenous PRG led to increased RhoA-GTP levels over basal, which was further significantly increased with co-expression of Gα_i1_-QL but not Gα_i1_-WT (Fig. 4B). These assays establish that Gα_i1_ activates the RhoGEF activity of PRG and the activation is dependent on the GTP-bound state of Gα_i1_.

### PRG-dependent SRE-Luc Activation is Specific to Gα_i/o_ Proteins

Activation of PRG by Gα_i1_ is surprising given that previous studies have indicated PRG regulation by Gα_12/13_ [22, 29]. We tested if activation of PRG in the SRE-Luc reporter assay extends to different families of Gα subunits. We co-expressed either WT or QL versions of different classes of Gα subunits, Gα_q_, Gα_12_, Gα_13_ as well as members of the Gα_i_ family subunits Gα_oA_, Gα_oB_ and Gα_z_ and Gα_i1_ with SRE-Luc, with or without PRG (Fig. S3A). Transfection of both WT and QL versions of Gα_q_, Gα_12_, Gα_13_ increased the luminescence signal compared to basal, likely due to activation of endogenous RhoGEFs in A293 cells. Under the conditions of this assay, Gα_q_, Gα_12_, Gα_13_ co-expression with PRG did not significantly increase PRG-dependent SRE-Luc activation. Increasing the expression of these Gα subunits increased basal SRE-Luc activity but did not synergize with PRG. The lack of effect of Gα_12_ or Gα_13_ in this assay is surprising given the well-accepted role of Gα_12/13_ in PRG regulation but is in accordance with a general lack of data demonstrating Gα_12/13_-dependent activation of PRG in cell-based co-transfection assays (see discussion).

### Gα_i1_ and Gα_i3_ Strongly Activate PRG, but Gα_i2_ is a Poor Activator

We tested Gα_i_ family member isoforms for their ability to activate PRG. Gα_oA_-QL, Gα_oB_-QL showed a small increase in PRG activation while Gα_z_ did not activate PRG at all (Fig. S3B). Gα_i_ isoforms Gα_i1_, Gα_i2_, and Gα_i3_ are highly homologous, with greater than 85% amino acid sequence identity [6]. Strikingly, Gα_i1_-QL and Gα_i3_-QL strongly activated PRG, while Gα_i2_-QL did not (Fig. 5A). Gα_i1_ and Gα_i2_ were expressed approximately equally (Fig. 5B), and Gα_i3_ expression could not be compared because the antibody does not recognize Gα_i3_. Gα_i_-QL versions of all three subtypes showed similar inhibition of Fsk-stimulated cAMP production in a parallel assay (Fig. 5C). To corroborate this finding, we co-expressed PRG with BioID2 tagged WT and QL versions of Gα_i1_, Gα_i2_, Gα_i3_, and BioID2-CaaX in A293 cells. The cells were labeled with biotin for 24 hr, followed by streptavidin affinity pull-down and western blotting for PRG. PRG was highly enriched in the BioID2-Gα_i1_-QL samples, followed by BioID2-Gα_i3_-QL and very low labeling by BioID2-Gα_i2_-QL. Effector selectivity amongst these three highly related Gα_i_ isoforms has not been previously reported.

**Figure 5.**
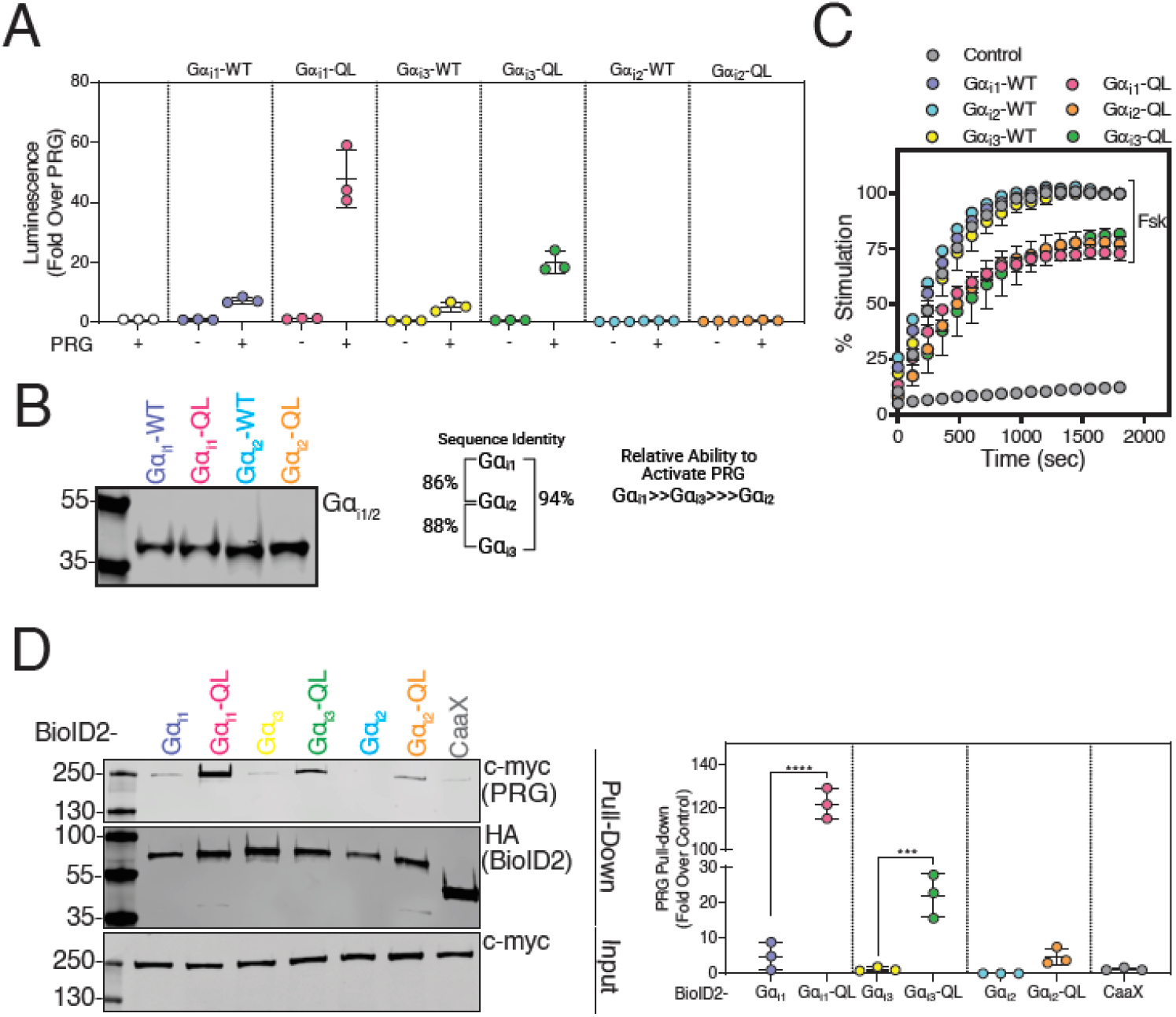
PRG Activation is Gα_i_ Isoform Specific. **(A)** Gα_i1_-QL and Gα_i3_-QL strongly activate PRG, but Gα_i2_ poorly activates PRG. cDNAs for each of the indicated Gα protein subunits were transfected with PRG and SRE-Luc and assayed as in Fig 4A. **(B)** Western blot showing relative expression of Gα_i1_ and Gα_i2_ constructs in A293 cells. **(C)** Gα_i1_, Gα_i2_, and Gα_i3_-QL equally inhibit cAMP accumulation. Cells were transfected with cAMP Glosensor™ and WT and QL versions of Gα_i1_, Gα_i2_, or Gα_i3_ for 24 hr. Luminescence was measured for 30 min (x-axis) after Forskolin (Fsk) stimulation and represented as % stimulation (y-axis) relative to the maximum signal in the respective WT group with 1 μM Fsk treatment. **(D)** Enrichment of PRG by BioID2-Gα_i1_-QL and BioID2-Gα_i3_-QL relative to BioID2-Gα_i2_-QL confirmed by biotin labeling coupled to pull-down and western blotting. A293 cells were transfected with indicated constructs, labeled in the presence of biotin for 24 hr, and subsequently pulled down using streptavidin. Left panel: Representative western blots of PRG isolated using streptavidin pull-down as well as expression of BioID2-Gα_i1_, Gα_i1_-QL and BioID2-CaaX. Right panel: Quantitation of three independent experiments normalized to total PRG. The data represent the mean ± SD of three separate experiments. (n=3, one-way ANOVA. ***P < 0.001, ****<0.0001).

### Gα_i1_-QL is Specific for PRG Mediated RhoA Activation and Does Not Require Gα_12/13_

As discussed above, previous reports show that Gα_12/13_ binds to PRG [22, 29]. To investigate if Gα_12/13_ is required for Gα_i1_-QL mediated PRG activation, we performed the SRE-Luc reporter assay in wild-type and Gα_12/13_ null A293 (ΔGα_12/13_) cells, generated by CRISPR/Cas9-mediated gene editing. Gα_i1_-QL co-transfected with PRG robustly increased reporter activity, and knockout of Gα_12/13_ did not affect PRG activation (Fig. S4A left panel). A western blot of endogenous Gα_13_ compared between wild-type and ΔGα_12/13_ A293 confirmed the Gα_13_ KO (Fig. S4A right panel). This demonstrates that Gα_i1_-GTP can activate RhoGEF activity of PRG in the absence of Gα_12_ or Gα_13_.

Next, we tested the ability of Gα_i1_-QL to activate other members of the DH-PH family of RhoGEFs in the SRE-Luc reporter assay (Fig. S4B left panel). We co-transfected Gα_i1_-QL with GFP-p115 RhoGEF (Lsc), GFP-LARG, GFP-AKAP13 (Proto-Lbc, ARHGEF13), and GFP-PRG. These GFP-fused proteins were active and produced significant basal activities when transfected at higher cDNA concentrations (data not shown). Gα_i1_-QL robustly increased SRE-Luc reporter activity when co-transfected with PRG. Statistically significant synergistic activation by Gα_i1_-QL was also observed for p115RhoGEF, although this was much lower as compared to PRG activation. No statistically significant activation of LARG or AKAP13 was observed. Western blotting demonstrated expression of the RhoGEFs although at different levels. (Fig. S4B right panel).

### The Formyl Peptide Receptor 1 (FPR1) Activates PRG via Gα_i1_

To determine whether PRG could be activated by Gα_i_ downstream of a Gi-coupled receptor, we assayed PRG activation using SRE-Luc in A293 cells stably expressing FPR1 (A293-FPR1). fMLF activated SRE-Luc reporter activity in a concentration-dependent fashion only when PRG and Gα_i1_-WT were co-transfected and not with PRG or Gα_i1_-WT alone (Fig. 6A). In contrast, fMLF did not increase reporter activity in cells co-transfected with Gα_i2_-WT and PRG (Fig. 6A), recapitulating the finding that Gα_i2_-QL poorly activates PRG. To further show that FPR1 activation of PRG depends on Gα_i_, cells expressing FPR1, Gα_i1_ and PRG were treated with PTX. As expected, PTX significantly inhibited the fMLF-dependent increase in PRG activation (Fig. 6B).

**Figure 6.**
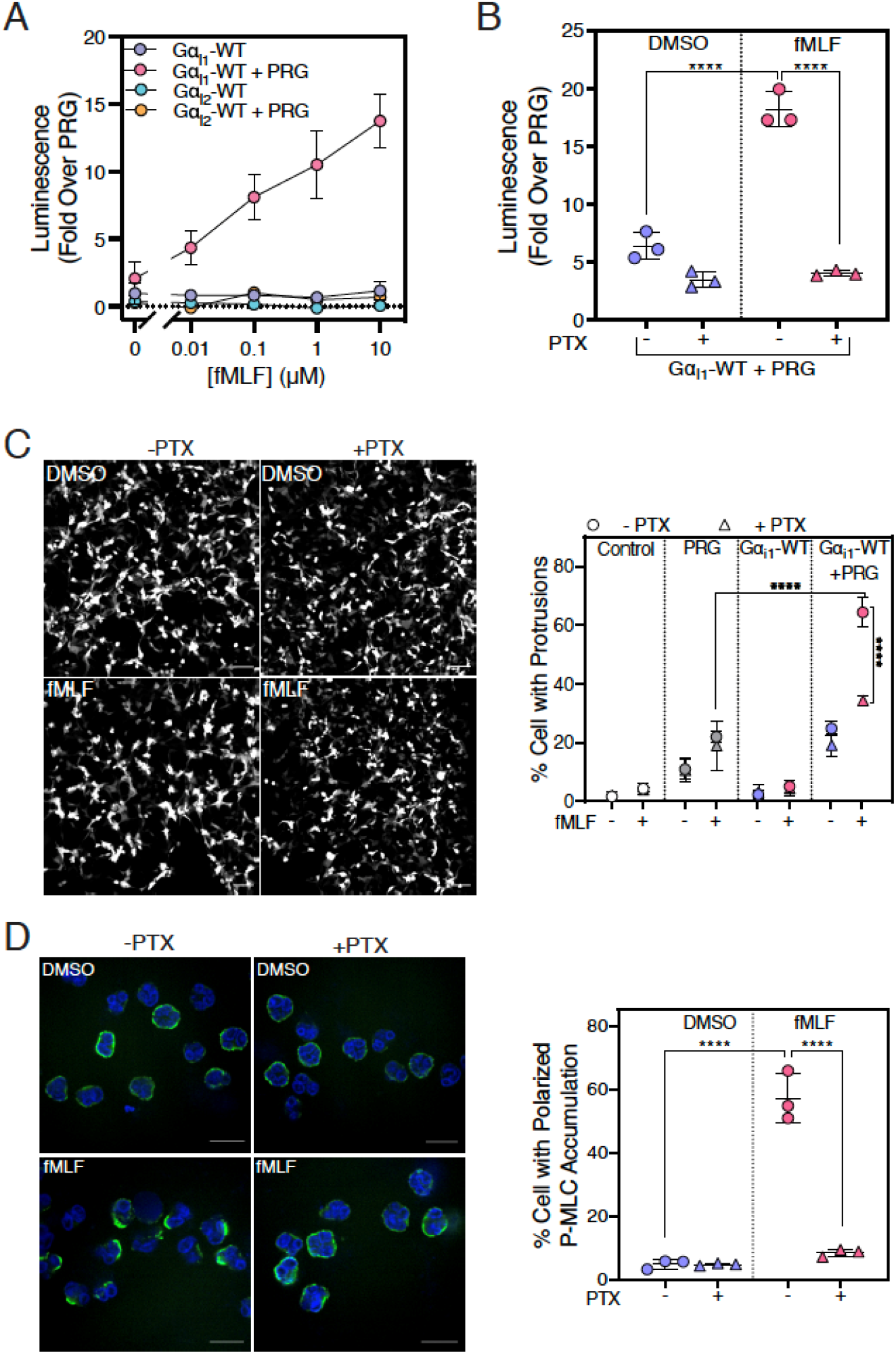
Activation of PRG Downstream of Gα_i_-coupled Receptor FPR1. **(A)** fMLF activates PRG in A293-FPR1 cells transfected with Gα_i1_-WT but not Gα_i2_-WT. A293-FPR1 cells were transfected with cDNAs encoding SRE-Luc, PRG, and Gα_i1_-WT or Gα_i2_-WT and subsequently incubated for 12 hr in serum free media containing the indicated concentrations of fMLF (0.01, 0.1, 1 and 10 μM) or DMSO. **(B)** PTX inhibits fMLF-mediated PRG activation in A293-FPR1 cells transfected with Gα_i1_-WT. The cells were transfected, and serum starved as described in 6A. Cells were treated with PTX (100 ng/mL) along with 10 μM fMLF stimulation for 12 hr. Luminescence was measured 10 min after the addition of the One-Glo™ reagent. **(C)** fMLF stimulation of A293-FPR1 cells transfected with PRG and Gα_i1_-WT increases the number of cells with dynamic protrusions (video S1-S4). A293-FPR1 cells were transfected with PRG, Gα_i1_-WT, and YFP for 36 hr. For PTX treatment, cells were treated with PTX (100 ng/mL), 24 hr after transfection, for 12 hr. Subsequently, the cells were stimulated with 100 nM fMLF, and live cell video microscopy was performed for 40 min. Left panel: Representative images (10×) of A293-FPR1 cells expressing PRG + Gα_i1_-WT and treated with fMLF or DMSO are shown. Scale bar, 100 μm. Right panel: Quantitative analysis of three independent experiments. For each experiment, >500 cells per condition were analyzed in a blinded manner and quantitated for the percentage of cells with dynamic protrusions. **(C)** fMLF stimulation increased the number of cells with polarized P-MLC accumulation. Human neutrophils were pretreated with or without PTX (500 ng/mL) for 2 hr and allowed to adhere to the fibronectin-coated surface for 15 min and stimulated with 10 nM fMLF for 5 min. Subsequently, the cells were fixed, permeabilized, stained using an anti-P-MLC antibody, DAPI, and imaged using confocal miscopy. Left panels: Three random fields were imaged, and representative images are shown. Right panel: Total number of cells and cells with asymmetric P-MLC localization were counted in a field and % cells with polarized P-MLC localization from three independent experiments were plotted (right panel). The data represent the mean ± SD of three separate experiments. (n=3, one-way ANOVA. ****P < 0.0001). Scale bar, 10 μm.

RhoA is a major regulator of cytoskeletal rearrangement and can induce peripheral protrusions [30, 31]. As an alternate measure of RhoA activation, we examined fMLF-stimulated dynamic protrusion formation in FPR1-A293 cells (Fig. 6C left panel, Videos S1-4). The percentage of cells that formed protrusions in cells transfected with PRG alone was slightly but significantly higher than in vector control cells and was unaffected by PTX treatment (Fig. 6C right panel). Co-expression of Gα_i1_-WT with PRG significantly increased the percentage of cells with dynamic protrusions only when cells were stimulated with fMLF, and the activation was strongly inhibited by PTX. These data together support the hypothesis that PRG RhoGEF activity can be activated downstream of Gi-coupled GPCRs through Gα_i1_.

### fMLF-dependent PRG Activation Requires Gα_i_ in Human Neutrophils

To understand the physiological relevance of this fundamentally new Gα_i_-dependent mechanism for Rho regulation, we examined the role of Gi signaling in human neutrophils. Previous studies have shown that of the multiple G protein-activated RhoGEFs expressed in neutrophils, PRG mediates Rho-dependent polarized accumulation of phosphorylated (at serine-19) -myosin light chain (P-MLC) at the trailing edge of migrating neutrophils [20, 21]. This has been proposed to result from FPR1-dependent Gα_13_ activation [20] in part because PTX treatment of HL60 cells, a neutrophil-like cell line, only partially inhibited fMLF-dependent Rho activation and asymmetric localization of P-MLC [20]. To determine if Gα_i_ activates endogenous PRG in human neutrophils, we examined polarized P-MLC staining in human neutrophils after stimulation with a physiologically relevant concentration of fMLF (10 nM). In DMSO treated neutrophils, P-MLC was uniformly distributed at the surface of cells. Stimulation with 10 nM fMLF promoted strong polarized P-MLC accumulation (Fig. 6D left). In cells pretreated with PTX, 10 nM fMLF failed to promote polarized accumulation of P-MLC (Fig. 6D). Since PRG is responsible for Rho-dependent P-MLC polarization, this finding indicates that at physiological concentrations of fMLF, PRG activation in human neutrophils is via a Gα_i_-dependent mechanism.

## Discussion

In this study, we used an unbiased approach to identify novel effectors of and functions for, Gα_i_ and focused on identifying proteins involved in chemoattractant-dependent cell migration. Previous studies have screened for novel Gi targets using yeast two-hybrid systems or IP- followed by MS [15–17]. The proximity labeling method used in this study has numerous advantages over these other methods allowing for detection of transient complex formation in the context of an intact cell. One potential drawback is that apart from detecting direct or indirect interactions, proximity-based methods can also identify the proteins that do not interact but are located within 20 nm, perhaps in the same cellular compartment. However, the ratiometric enrichment strategy employed here comparing constitutively active Gα_i_ to inactive Gα_i_ largely circumvents this issue. GTP binding to Gα subunits lead to well-defined conformational changes that drive new protein-protein interactions. In principle, selective enrichment in the GTP-bound state could result from a few processes: 1) GTP-selective protein-protein interactions causing an increase in proximity, 2) GTP driven changes in Gα_i_ subunit compartmentation within the membrane or cell, or 3) Gα_i_-GTP driven changes in selective protein expression. There is some evidence that Gα_i_ activation leads to changes in compartmentation within the PM but not for larger-scale changes in subcellular distribution [32]. There is overwhelming evidence that G protein α subunit activation results in conformational changes that drive new protein interactions [3]. G protein targets may not be easily identified by traditional methods due to their characteristically low abundance, cell context specificity, and often transient interactions. Thus, proximity labeling, with proper controls, is a novel approach for investigation of Gα_i_ subunit interactions in intact cells and may represent a general approach for identification of signaling partners and networks downstream of G proteins.

Multiple known Gα_i_ binding partners were identified in the proximity labeling MS experiments. Gβ subunits, Gγ subunits, and GPSM1 (AGS3) [33] were enriched in inactive Gα_i1_ samples, whereas GPRIN1[26], Rasa3 [16], and GIV [34] were enriched in Gα_i1_-QL samples. Many AC isoforms were identified but somewhat surprisingly were equally enriched in both Gα_i1_ and Gα_i1_-QL samples. It has been suggested that Gα_i_-AC complexes can be formed regardless of the activation state of Gi, and that AC activation results from conformational rearrangement of the prebound G protein heterotrimer [35–37]. GPCRs were detected, but the overall labeling efficiency was low for many of them, leading to lower confidence in identification for some of the receptors (Table S1), and most were not significantly enriched by either Gα_i1_ or Gα_i1_QL. This could be because of their low receptor abundance or lack of preference for the inactive receptors for Gα_i1_ or Gα_i1_QL. Overall, many bona fide Gα_i_ targets were identified in the MS screen, validating the method, suggesting that novel Gα_i_ effectors are likely to be identified.

Proteins enriched in the Gα_i_-QL samples were grouped into unique functional categories, including classes related to cell migration (Fig 2C, Table S2). The fact that multiple proteins were identified suggests that the role of Gα_i_ in cell migration may involve a network of protein-protein interactions similar to the role of Gβγ in cell migration. These potential interactions remain to be independently investigated, but many are likely “true” interaction partners. Indeed, the follow-up biotinylation validation assays with “randomly” selected, BioID2-Gα_i1_-QL enriched proteins (Fig. 2E) support the idea that many of these proteins selectively interact with activated Gα_i_. In a process with the complexity of chemoattractant-dependent cell migration, Gα_i_ is likely to play more than one role.

From the collection of proteins identified in this study, we focused on PRG, a RhoGEF with well-established roles in cell migration, and demonstrated that Gα_i1_ strongly activates PRG in a GTP and concentration-dependent manner in intact cells. We also demonstrated Gα_i_-dependent PRG regulation downstream of the Gi-coupled chemoattractant receptor, FPR1, in A293 cells. In differentiated HL60 neutrophil-like cells, activation of FPR1 leads to PRG and Rho-dependent accumulation of P-MLC at the trailing edge, where it is responsible for tail retraction as the cell moves forward [20, 21]. This has been proposed to be mediated by Gα_13_. In our experiments, stimulation of primary human neutrophils with a physiological concentration fMLF promoted polarized P-MLC accumulation that was completely inhibited by PTX, confirming this phenotype to be dependent on Gα_i_ signaling. In the previous work performed with 100 nM fMLF, PTX did have some effect on P-MLC polarization, and fMLF dependent RhoA-GTP was partially inhibited by PTX [20]. Some differences between this previous study and our work showing a complete Gα_i_-dependence of P-MLC polarization are the use of primary neutrophils and the use of physiological concentrations of fMLF (10 nM). The data suggest that at low physiological concentrations of chemoattractant, Gα_i_ regulation of PRG is critical for tail retraction during cell migration, and we propose that Gα_13_ may perform a more dominant role in migrating neutrophils at higher concentrations of chemotactic ligand.

It is well accepted that Gα_13_ activates PRG and other RH-RhoGEFs, and regulation by Gα_i_ has not been previously documented. We did not observe activation of PRG by Gα_12_ or Gα_13_ in our assays. Most of the published evidence convincingly yet indirectly shows that PRG activity is regulated downstream of Gα_12/13_ [22, 38]. Despite strong data demonstrating G_13_ binding to PRG and clear regulation of PRG by G_12/13_ coupled GRCRs in physiological settings [39], demonstration of Gα_12/13_-dependent regulation of PRG in cell-based assays similar to those used in our studies are limited [40, 41]. The reasons for these discrepancies are unclear. A recent report demonstrated Gα_s_ mediated activation of Cdc42 via the DH-PH domain of PRG [42]. Thus, there is precedence for regulation of PRG by G proteins other than G_12/13_.

From our studies, we cannot conclude whether the mechanism for Gα_i_-dependent regulation of PDZ-RhoGEF involves direct protein-protein interactions or if a higher-order complex is involved. The strong stimulation of RhoGEF activity of PRG by Gα_i_ suggests direct interactions, but further *in vitro* reconstitution experiments with purified components will be required to conclusively demonstrate direct PRG regulation by Gα_i_.

Of the three highly homologous Gα_i_ isoforms, Gα_i1_, Gα_i2_, and Gα_i3_; Gα_i1_ and Gα_i3_ activated PRG, but surprisingly Gα_i2_ was a poor activator [13]. Gα_i1_-QL shares 86% amino acid identity with Gα_i2_ [43, 44], and the three Gα_i_ isoforms inhibit AC with similar potency and efficacy [6]. These three Gα_i_ isoforms have been studied for nearly three decades [43], and no molecular differences with respect to effector regulation have been demonstrated. Mouse neutrophils equally express Gα_i2_ and Gα_i3_, and neutrophils from Gα_i3_ KO mice showed reduced ability to migrate toward a chemotactic stimulus, whereas Gα_i2_ KO resulted in loss of their ability to arrest [45, 46]. Therefore, this divergent role could be attributed to differential effector regulation by different Gα_i_ subtypes.

PRG is known to regulate a variety of biological processes, including neurite retraction [47], cell migration, and proliferation of mouse embryonic fibroblasts [48]. Thus, these findings have broad implications for signaling by Gi-coupled receptors. Overall, identification of signaling pathways and networks regulated by G_i_-coupled GPCRs has the potential to significantly impact our understanding of the biology regulated by these ubiquitous and pharmacologically important receptors. Identification of proteins from a range of functional families indicates that Gi proteins likely play a central role in multiple biological processes via signaling networks, far beyond just adenylate cyclase inhibition. Fuller characterization of individual candidate interactors or networks is warranted to validate and understand the roles of these interactions in Gi-coupled GPCR biology. Ultimately, discovery of new GPCR biology will lead to a greater understanding of disease pathologies, identification of novel therapeutic targets, and development of innovative therapeutic strategies.

## Materials and Methods

### Plasmid cDNA Constructs

BioID2 fused N-terminally with c-myc tag and C-terminally with mVenus followed by CaaX PM targeting motif (KKKKKKSKTKCVIM, derived from the C terminus of KRas), was a gift from Dr. Sundeep Malik, University of Rochester. Final clone: c-myc-BioID2-mVenus-CaaX. C-terminally c-myc tagged full-length PRG cDNA construct in mammalian expression vector was a gift from Dr. John Tesmer, Purdue University. A293-FPR1 stable cell lines were generated by Dr. Jesi To, University of Michigan, and A293-Gα_12/13_ CRISPR cells were a gift from Dr. Graeme Milligan, University of Glasgow, UK. Following plasmids were obtained from Addgene. MCS-BioID2-HA (Kyle Roux, Plasmid #74224) [19], pCDNA3-HA PSPC1(Yuh-Shan Jou, Plasmid #101764) [49], FLAG-p54 (Benjamin Blencowe, Plasmid # 35379) [50], pEGFP-ATF6-(S1P-) (Ron Prywes, Plasmid #32956) [51], GFP-nArgBP2 (Guoping Feng, Plasmid # 74514) [52], GFP-Golgin-84-TEV (Ayano Satoh, # 42108), mEmerald-Parvin-C-14 (Michael Davidson, Plasmid # 54214), EGFP-Vimentin-7 (Michael Davidson, Plasmid # 56439), pGFP-Cortactin (Kenneth Yamada, Plasmid # 50728). All Gα clones in pcDNA3.1+ were obtained from the cDNA Resource Center. Gα_i1_-FLAG-APEX2, Gα_i1_-QL-FLAG-APEX2, Lyn-FLAG-APEX2, and EGFP-BioID2-HA-CaaX were synthesized by GenScript. The sequences of the clones are available upon request.

### Design and Cloning of cDNA Constructs

BioID2-HA was inserted between 121-Alanine and 122-Glutamic acid of human Gα_i1_-WT and Gα_i1_-QL with a linker (residues- SGGGGS) flanking BioID2-HA on either side. Final clone: (Gα_i1_(1-121)-Linker-BioID2-HA-Linker-Gα_i1_(122-355). GFP-PRG was generated from PRG amplification from FL-c-myc-PRG and insertion into the pEGFP-N1 vector.

### Cell Culture

A293 and HT1080 cells were obtained from American Type Culture Collection (ATCC). A293, A293-FPR1, and HT1080 cells were grown in DMEM (10013CV, Corning) supplemented with 10% fetal bovine serum (FBS) (10437028, Gibco) and 100 units of penicillin/streptomycin (P/S) (15140122, Gibco) at 37 °C with 5% CO_2_. Media was supplemented with 100 μg/mL Geneticin (G418) (G8168, Sigma) to select A293-FPR1 cells. Trypsin-EDTA (25200056, Gibco) was used for cell passaging.

### Reagents

The following primary and secondary antibodies were utilized: Gα_i1/2_ (anti-sera) [53], PDZ-RhoGEF (ab110059, abcam), HA (3724, Cell Signaling), FLAG (F1804, Sigma), P-MLC (3671, Cell Signaling), c-myc (13-2500, Invitrogen), Streptavidin-IRDye800 (925-32230, LI-COR), GFP (A11122, Invitrogen). Primary antibodies were made in 3% BSA and 0.1% Sodium azide and the blots were incubated in primary antibody overnight at 4 °C except 1 hr incubation at RT for streptavidin-IRDye800. Secondary antibody goat anti-rabbit DyLight™ 800 (SA535571, Invitrogen), goat anti-mouse IRDye 800CW (926-32210, LICOR) at 1:10,000 dilution and goat anti-rabbit Alexa Fluor 488 (A11034, Invitrogen) at 1:1000 dilution.

### Proximity-Based Labeling using BioID2

#### Small-scale total protein biotinylation and western detection

A293 cells were plated in a 6-well plate at a density of 0.35 × 10^6^ cells per well. 24 hr after plating, media was replaced with 2 mL DMEM supplemented with 50 μM Biotin (B4501, Sigma) (prepared as previously described [54] and 10% FBS. The cells were then transfected with 1 μg of BioID2 clone (BioID2-Gα_i1_, BioID2-Gα_i1_-QL or BioID2-CaaX) and 100 ng of yellow fluorescent protein (YFP) cDNAs in each well using 1:3 DNA: Lipofectamine 2000 (11668019, Invitrogen) ratio. 24 hr after concurrent transfection and biotin labeling, 300 μL 1× Laemmli buffer was added per well, and the lysates were collected, boiled for 10 min at 95 °C, 40 μL of was resolved on 4-20% Mini-protean TGX™ Gel (4561094, Bio-Rad) and detected by western blot. Anti-HA (1:2000), anti-PDZ-RhoGEF (1:1000), anti-c-myc (1:2000), Streptavidin-IRDye800 (1:3000) were utilized.

#### Protein biotinylation, pull-down, and western detection

A293 cells were plated in a 10 cm dish at a density of 2.0 × 10^6^ cells per dish. The next day, media was replaced with 10 mL DMEM supplemented with 50 μM Biotin and 10% FBS. Thereafter, the cells were transfected with 3 μg of BioID2-Gα_i1_, BioID2-Gα_i1_-QL, or BioID2-CaaX and 3 μg of protein of interest (HA-PSPC1, FLAG-p54-HA, GFP-ATF6, GFP-ArgBP2, Golgin A5-GFP, Parvin-GFP, Vimentin-GFP, Cortactin-GFP) in each dish using 1:3 DNA: Lipofectamine 2000 ratio. 24 hr after transfection and labeling, the cells were harvested by centrifugation at 4000× g for 10 min and lysed in 500 μL ice-cold lysis buffer (modRIPA buffer: 50 mM Tris, 150 mM NaCl, 0.1% SDS, 0.5% Sodium deoxycholate, 1% Triton X-100, final pH 7.5) supplemented with 1×protease inhibitor (PI) cocktail (P8849, Sigma), 1 mM Phenylmethylsulfonyl fluoride (PMSF) (786-055, G-Biosciences) for 10 min on ice. The lysates were further incubated with 125 units of Benzonase (E1014-25KU, Sigma) in an end-over-end rotator at 4 °C for 20 min. 0.3% SDS was added to lysates and incubated for an additional 10 min at 4 °C. Lysates were centrifuged at 15000× g for 10 min, and the supernatant was transferred to fresh tubes and total protein concentration was equalized using Pierce 660-nm protein assay reagent (22660, Thermo Fisher Scientific). 5% of equalized lysates were taken out before pull-down to analyze the biotinylation of inputs by western blot analyses. The remaining lysates were incubated with 100 μL Pierce™ streptavidin magnetic beads slurry (88817, Thermo Fisher Scientific) per sample in an end-over-end rotator at 4 °C for 18 hr to capture biotinylated proteins. Following streptavidin pull-down, beads were washed twice with ice-cold modRIPA, and once with ice-cold 1× PBS. 30 μL 1× Laemmli buffer was added to the beads, boiled for 10 min at 95 °C, and the supernatant was loaded on the SDS-PAGE followed by western blot analyses. HA (1:2000), PDZ-RhoGEF (1:1000), c-myc (1:2000), mCherry (1:1000) were utilized.

#### Large scale protein biotinylation and pull-down

Low passage HT1080 cells (passage number up to 15) were used for proximity labeling experiments. HT1080 cells were plated into 175 cm^2^ flasks at a density of 5.5 × 10^6^ cells per flask. The next day, media was replaced with 35 mL DMEM containing 50 μM biotin and 10% FBS. Subsequently, the cells were transfected with 8 μg of BioID2 and 4 μg of YFP cDNAs in each flask. 0.6 μL of Viromer® Red (VR-01LB-00, Lipocalyx, Germany) reagent was used per 2 μg of cDNA for transfection, resulting in ~80-85% transfection efficiency. 24 hr after labeling and transfection, the labeling medium was decanted, cells were washed twice with 1×PBS, and harvested at 4000× g for 10 min. This step was repeated twice using 1×PBS to recover the maximum number of cells. The supernatant was aspirated, and pellets were snap-frozen and stored at −80°C until further use.

All stock solutions used for streptavidin pull-down were freshly prepared, except lysis buffer. Low protein binding tubes (022431081, Eppendorf) were used for sample preparation. Frozen pellets were lysed in 1 mL of ice-cold lysis solution (composition described above) for 10 min on ice, incubated with 125 units of Benzonase with end-over-end rotation at 4 °C for 20 min. 0.3% SDS was added to lysates and incubated for another 10 min at 4 °C. Lysates were centrifuged at 15,000×g for 15 min, the supernatant was transferred to fresh tubes, and total protein concentration was equalized using Pierce 660-nm protein assay reagent. 5% of equalized lysates were saved before pull-down to analyze the biotinylation of inputs by western blot analysis. The remaining equalized lysates were incubated with 500 μL Pierce™ streptavidin magnetic beads slurry per sample, in an end-over-end overnight 4°C for 18 hr. Subsequently, the beads were washed twice with modRIPA, once with four different solutions: 1 M KCl, 0.1 M Na_2_CO_3_, 2% SDS (made in 50 mM Tris pH 7.5), and 2 M Urea (made in 10 mM Tris pH 8.0). Finally, the beads were washed twice with 1× PBS and were snap-frozen and stored at −80 °C until further processing for mass spectrometry.

### Protein Digestion and TMT Labeling

On-bead digestion followed by LC-MS/MS analysis was performed at the mass spectrometry-based Proteomics Resource Facility of the Department of Pathology at the University of Michigan. Samples were reduced (10 mM DTT in 0.1 M TEAB at 45°C for 30 min), alkylated (55 mM 2-chloroacetamide at room temperature (RT) for 30 min in the dark, and subsequently digested using 1:25 trypsin (V5113, Promega): protein at 37° C with constant mixing using a thermomixer. 0.2% TFA was added to stop the proteolysis, and peptides were desalted using a Sep-Pak C18 cartridge (WAT036945, Waters Corp). The desalted peptides were dried in a Vacufuge and reconstituted in 100 μL of 0.1 M TEAB. A TMT10plex™ isobaric labeling kit (0090110, Thermo Fisher Scientific) was used to label each sample per manufacturer’s protocol. The samples were labeled with TMT 10-plex reagents at RT for 1 hr. The reaction was quenched by adding 8 μL of 5% hydroxylamine for 15 min, combined, and subsequently dried. An offline fractionation of the combined sample into 6 fractions was performed using a high pH reversed-phase peptide fractionation kit, as per manufacturer’s protocol (84868, Pierce). Fractions were dried and reconstituted in 12 μL of 0.1% formic acid/2% acetonitrile for LC-MS/MS analysis. Sample-to-TMT channel information is provided below.

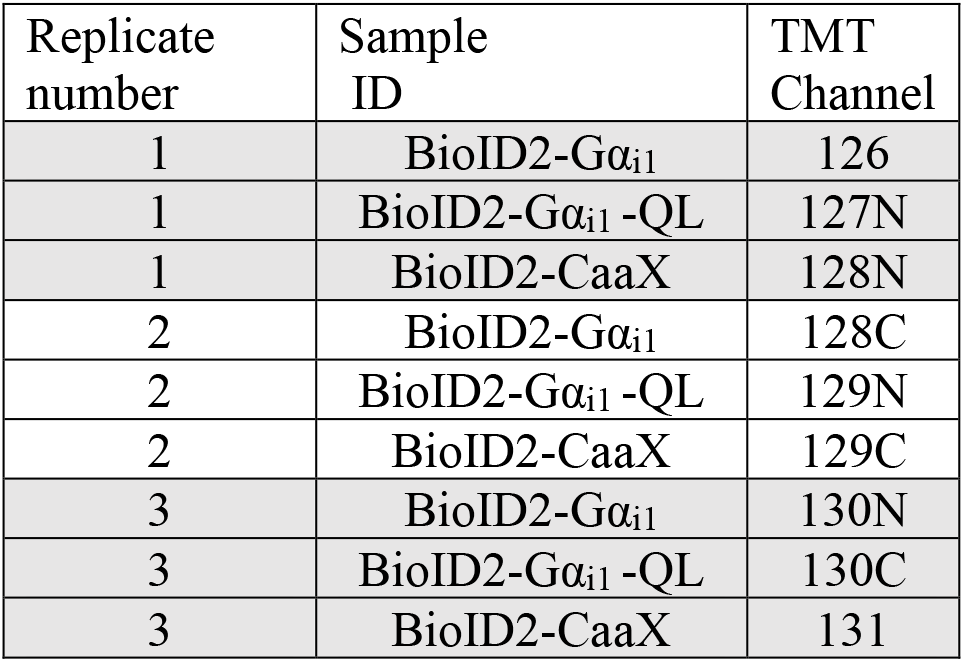

### Liquid Chromatography-Mass Spectrometry Analysis

An Orbitrap Fusion (Thermo Fisher Scientific) and RSLC Ultimate 3000 nano-UPLC (Dionex) was used to acquire the data. To achieve superior quantitation accuracy, we employed multinotch-MS3 [55]. 2 μL of each fraction was resolved on a nano-capillary reverse phase column (PepMap RSLC C18 column, 75 μm i.d. × 50 cm; Thermo Scientific) at the flow rate of 300 nL/min using 0.1% formic acid/acetonitrile gradient system (2-22% acetonitrile in 110 min;22-40% acetonitrile in 25 min; 6 min wash at 90% followed by 25 min re-equilibration) and directly sprayed onto the Orbitrap Fusion using EasySpray source (Thermo Fisher Scientific). Mass spectrometer was set to collect one MS1 scan (Orbitrap; 120K resolution; AGC target 2×10^5^; max IT 50 ms) followed by data-dependent, “Top Speed” (3 seconds) MS2 scans (collision-induced dissociation; ion trap; NCD 35; AGC 5×10^3^; max IT 100 ms). For multinotch-MS3, the top 10 precursors from each MS2 were fragmented by HCD followed by Orbitrap analysis (NCE 55; 60K resolution; AGC 5×10^4^; max IT 120 ms, 100-500 m/z scan range).

Proteome Discoverer (v2.4; Thermo Fisher) was used for data analysis. Tandem MS spectra were searched against SwissProt human protein database using the following search parameters: MS1 and MS2 tolerance were set to 10 ppm and 0.6 Da, respectively; carbamidomethylation of cysteines (57.02146 Da) and TMT labeling of lysine and N-termini of peptides (229.16293 Da) were considered static modifications; oxidation of methionine (15.9949 Da) and deamidation of asparagine and glutamine (0.98401 Da) were considered variable. Proteins and peptides that passed ≤1% false discovery rate threshold were retrained for subsequent analysis. Quantitation was performed using TMT reporter ion in MS3 spectra with an average signal-to-noise ratio of 10 and <50% isolation interference.

### Normalization and Sorting Criteria

Only a small fraction of all the proteins labeled by Gα_i1_ BioID2 are expected to have increased enrichment in BioID2-Gα_i1_-QL samples relative to the BioID2-Gα_i1_ samples. Most of the proteins are expected to be equally enriched across samples as the majority of the labeling is based on proximity rather than Gα_i1_-QL-specific interactions. Therefore, to quantitatively compare the samples across groups, we summed the total TMT signal for each sample to obtain a normalization factor used to normalize the values for each protein across experimental groups.

Normalized abundance ratio and p-values were used for the subsequent analysis. Proteins constituting the active Gα_i1_ interactome fulfilled all the following criteria: PSM>5, Abundance ratio BioID2-Gα_i1_/BioID2-CaaX ≥ 0.8 and BioID2-Gα_i1_-QL/ BioID2-Gα_i1_ ≥ 1.3, Abundance ratio P-value BioID2-Gα_i1_-QL/BioID2-CaaX <0.05.

### Gene ontology analysis

Gene ontology (Go) analysis was performed using the DAVID Bioinformatics resource at https://david.ncifcrf.gov. Proteins selected based on the criteria in figure 2 C were submitted based on gene identifiers to the analysis server and analyzed by functional annotation clustering.

### Immunofluorescence Staining

A293 cells (2 × 10^4^ cells/well) were plated on a poly-D-lysine coated 8-well chamber μ-slide (80826, Ibidi) and transfected with BioID2 clones (200 ng cDNA/well) the following day using 1:3 DNA: Lipofectamine 2000 ratio. 24 hr post-transfection, cells were fixed with 4% (w/v) paraformaldehyde (PFA) (15710, Electron microscopy sciences) for 10 min at RT and washed with phosphate-buffered saline (PBS, BP3994, Fisher). Subsequently, cells were blocked and permeabilized with 10% normal goat serum in 1× PBS containing 0.1% (v/v) Triton X100 (1× PBS-T) for 1 hr at RT. Primary anti-HA antibody HA (3724, Cell Signaling) was used at 1:500 dilution in 2% goat serum in 1×PBS-T overnight at 4°C. The next day, cells were washed three times with 1× PBS-T and incubated with secondary antibody (goat anti-rabbit Alexa Fluor 488) at a dilution of 1:1000 in 1× PBS-T for 1 hr at RT. The nuclei were stained with DAPI for 15 min and washed once with 1× PBS-T and 1× PBS. Cells were imaged on a LEICA DMi8 microscope in confocal mode with a 63× oil lens using 405 nm excitation for DAPI and 488 nm for Alexa Fluor 488 secondary antibody. Acquisition parameters were kept constant for all the conditions of an experiment.

### Glosensor c-AMP Reporter Assay

A293 cells (4 × 10^4^ cells/well) were plated per well in a 96-well plate (655983, Greiner). The following amounts of DNA were used per well: 50 ng of pGloSensor™ −20F cAMP Plasmid (E1171, Promega), 125 ng of untagged Gα_i1_-WT, Gα_i1_-QL or BioID2 fused Gα_i_ clones or empty vector (control, pCDNA3.1+). Reverse transfection was performed using 1:3 DNA: Lipofectamine 2000 ratio. 24 hr after transfection, cells were washed once with 1× PBS and 75 μL of 2 mM D-luciferin (LUCK-1G, Goldbio) in Leibovitz’s L-15 medium (21083-027, Gibco) was added for 2 hr at 37 °C, 5% CO_2_ incubator. Cells were treated with vehicle or 1 μM forskolin (Fsk) (11018, Cayman Chemicals), and luminescence was measured using a Varioskan™ LUX multimode microplate reader for 30 min.

### Proximity Ligation Assay (PLA)

*In situ* PLAs were performed using Duolink™ Kit (DUO92101, Sigma) as per manufacturer’s protocol with some modifications. 2 × 10^4^ cells were plated on 14 mm coverslips in a 35 mm dish (D11030H, Matsunami) and the following day, 100ng of Gα_i1_-FLAG-APEX2, Gα_i1_-QL-FLAG-APEX2 or Lyn-FLAG-APEX2 with 25 ng EGFP-PRG were transfected and the cDNAs were allowed to express for 48 hr with media change after 24 hr. The cells were then washed twice with 1× PBS and fixed with 4% PFA, and 4% sucrose made in 1× PBS for 10 min in the dark at RT. The cells were permeabilized and blocked using freshly prepared 5% goat serum, 1% BSA, 0.2% Triton X100 in TBS. Subsequently, rabbit anti-GFP (1:750) (A11122, Invitrogen) and mouse anti-FLAG-(1:750) (F1804, Sigma) antibodies were diluted in Duolink^®^ antibody diluent and incubated overnight at 4 °C in a humidified chamber. A total 40 μL reaction mixture including PLA probe binding, ligation, amplification steps in a humidified chamber. The dilution factors for all the reagents were kept used as per manufacturer’s instructions. 2 mL of either buffer-A or B were used per wash as directed in the manual. After final washes, all the aqueous media was removed, 80 μL of mounting media was added to the cells. Random fields were imaged on the LEICA DMi8 microscope in confocal mode with a 63× oil lens, using 405 nm excitation for DAPI, 488 nm for GFP-PRG, and 568 nm for PLA dots. Acquisition parameters were kept constant for all the conditions of an experiment. The intensity of PLA dots and GFP-PRG was measured for ≥ 100 cells per condition, using ImageJ, and represented on the X and Y-axis, respectively.

### Luciferase Reporter Assay

A293 cells were plated, 4 × 10^4^ cells per well, in a 96-well plate (655983, Greiner). The following amounts of DNA were used per well: 25 ng of SRE luciferase reporter (E134A, Promega), 2.5 ng of c-myc-PRG, 0.75, 2.5.7.5,25,75,125 ng Gα_i1_ or Gα_i1_-QL. pcDNA3.1+ empty vector was used to keep the total amount of DNA constant in each well. Transfection was performed using 1:3 DNA: Lipofectamine 2000 (Invitrogen) ratio. Reverse transfection was performed, meaning cells were plated and transfected at the same time. 12 hr after transfection, media was replaced with 75 μL of serum-free media for another 12 hr, and 75 μL (1:1 volume) of One-Glo™ reagent (E6110, Promega) was added to each well, incubated for 10 min at RT. The luminescence signal was measured using Varioskan^TM^LUX multimode microplate reader (Thermo Scientific^TM^). A293-FPR1 cells were transfected and treated with 100 ng/mL PTX (P7208, Sigma) for 12 hr and subsequently, fMLF (0.01, 0.1, 1, 10 μM) was added in serum-free media with or without PTX, for the next 12 hr.

### Rhotekin Pull-Down Assay

Active RhoA levels were measured using the RhoA Pull-down Activation Assay Biochem Kit (BK036-S, Cytoskeleton Inc.) using GST Rhotekin beads. The levels of the GTP-RhoA associated with GST-Rhotekin-RBD were quantified by western blot analysis. Briefly, A293 cells were plated in a 6-well plate at a density of 3.5 x 10^5^ cells per well and transfected with 1 μg Gα_i1_, 100 ng c-myc-PRG and, 250 ng RhoA-HA per well using 1:3 DNA: Lipofectamine 2000 ratio. 20 hr after transfection, cells were cultured in serum-free media for 4 hr. Cells in each well were then lysed with 300 μL of RhoA lysis buffer with 1×PI (included with the kit), and lysates were equalized for total protein amount. Samples from two wells were pooled for each experimental group (total 600 μL, ~600 μg protein per experimental group). The lysates were incubated with 50 μg of GST-Rhotekin bound beads in an end-over-end rotator for 1 hr at 4 °C. Beads were washed twice with wash buffer (included with the kit), eluted in 40 μL 1× Laemmli sample buffer, and analyzed by western blot using an anti-HA (1:2000), anti-c-myc antibody (1:2000), and anti-Gα_i1/2_ antisera (1:3000). Band intensities were quantified using Image Studio Lite (version 5.2)

### FPR1-A293 Cell Protrusions Assay

FPR-1-A293 cells (2 × 10^4^ /well) were plated in an 8-well chamber slide with poly-D-lysine. The following plasmids were transfected per well: 100 ng YFP, 4 ng PRG, 125 ng Gα_i1_, or empty vector to bring the total amount of cDNA per well equal. Plasmids were transfected using 1:3 DNA: Lipofectamine 2000 ratio. 24 hr after transfection, the media was changed to fresh media, with or without PTX (100ng/mL) (P7208, Sigma) for 24 hr more. Subsequently, the cells were washed once with 1× PBS and placed in HBSS + HEPES (10mM) pH 7.3. The cells were imaged every 20 seconds for 40 min, and formyl-methionyl-leucyl-phenylalanine fMLF (F3506, Sigma) or vehicle was added 5 min after the video initiation. The videos were taken at 10× magnification on the LEICA DMi8 microscope using a 488 nm excitation filter. To quantify the % cells with protrusions, the total number of cells in first frame of a video were counted in ImageJ by ‘analyze particles’ option, and cell with the protrusions were manually counted from the videos, in a blinded manner.

### Human Neutrophil Isolation

Human peripheral blood was obtained from the Platelet Pharmacology and Physiology Core at the University of Michigan. The core maintains a blanket IRB for basic science studies, which doesn’t require HIPAA information, and enrolls healthy subjects that follow the protection of human subject standards. De-identified samples were used in the study.

Neutrophils were isolated from human peripheral blood as described previously [56]. Freshly isolated blood was carefully layered on top of 1-step polymorphs (AN221725, Accurate chemicals and scientific corporation) (1:1 Blood and Polymorphs) and centrifuged at 1000× g for 45 min and buffy coat was transferred to fresh tubes. Red blood cells were lysed using 0.1× PBS hypotonic solution for 45 sec, and immediately 4× PBS was added. The tubes were centrifuged at 400× g for 10 min, and pelleted cells were resuspended in modified Hanks’ balanced salt solution (mHBSS). Neutrophil preparations were at least 95% pure, as confirmed by nuclear morphology.

### Immunostaining of Human Neutrophils

Each well of an 8-well chamber μ-slide was coated with 5 μg of fibronectin (F1141, Sigma) overnight at 4°C. Freshly isolated human neutrophils were preincubated with either vehicle or 500 ng/mL PTX for 2 hr at 37 °C with gentle rotation before plating on the fibronectin-coated wells. 2 × 10^5^ cells per well were allowed to adhere to the surface for 15 min at 37°C in 5% CO_2_. The cells were stimulated with vehicle or fMLF (10 nM) for 5 min at 37 °C in 5% CO_2_ incubator and then fixed with 4% PFA and 5% sucrose in ddH_2_O for 15 min at RT, and blocked using 10% goat serum, 3% Fatty acid-free BSA, 0.05% Saponin in 0.2% PBST for 1 hr at RT. Subsequently, the cells were incubated with 1:100 P-MLC primary antibody prepared in 2% goat serum, 0.05% saponin in 0.1% PBST. The following day, the cells were washed with 0.05% saponin in 0.1% PBST for 10 min, three times, and incubated with anti-rabbit 488 secondary for 1 hr at RT and subsequently stained with DAPI. The cells were imaged in confocal mode with a 63× oil lens, and acquisition parameters were kept constant for all the experimental conditions. Three random fields were acquired per experiment, and images from three independent experiments were analyzed by counting the number of cells with asymmetric P-MLC staining and the total cells to determine % cells with asymmetric P-MLC distribution. Representative images were captured with a 100× oil lens.

### Western Blotting

Samples were resolved on 4-20% Mini-protean TGX™ Gels (4561094, Bio-Rad), were transferred to a nitrocellulose membrane (66485, Pall Corporation), and stained with Ponceau S (141194, Sigma). Membranes were blocked with 5% non-fat milk powder in TBST (0.1% Tween-20 in Tris-buffered saline) at RT for 1 hr with constant shaking. Membranes were probed with primary antibodies for 1 or 2 hr at RT or overnight at 4 °C. The membranes were washed with TBST, incubating with secondary antibody for 1 hr at RT, washed with TBST, and imaged using an Odyssey Infrared Imaging System (Li-Cor Biosciences).

### Statistical Analysis

All the experiments were performed at least three times, except Figure-2E, which was repeated twice. Data shown are expressed as mean ± SD or as one representative experiment. Statistical significance between various conditions was assessed by determining P values using the Student’s t-test or one-way ANOVA. Western blot images were scanned using Licor and quantified using Image Studio Lite (Version 5.2). All data were analyzed using GraphPad Prism 7.0 (GraphPad; La Jolla, CA), and schematic representations of the figures were created with BioRender.com and Adobe illustrator.

## Supplementary Materials

1. **Supplemental Figures:** Fig. S1. Characterization of BioID2 Fused Gα_i1_ and Gα_i1_-QL. Fig. S2. Heat Map of Proteins Identified as Enriched in BioID2-QL Samples Based on the Criteria in Fig. 2C Fig S3. Specificity of PRG Activation by Different G Protein Subunit Family Members in the SRE-luciferase Assay. Fig. S4. Gα_i1_-QL is Specific to PRG Relative to Other RhoGEFs and Does Not Require Gα_12/13_.
2. **Supplemental Tables:** Table S1. List of receptors identified by mass spectrometry. Table S2. Final 104 proteins. Table S3. Functional classification through Gene ontology analysis.
3. **Supplemental Videos:** Video S1. Protrusion dynamics: Gα_i1_+PRG+DMSO Video S2. Protrusion dynamics: Gα_i1_+PRG+FMLF Video S3. Protrusion dynamics: Gα_i1_+PRG+PTX+DMSO Video S4. Protrusion dynamics: Gα_i1_+PRG+PTX+FMLF

## Abbreviations

AC: Adenylate Cyclase
cAMP: 3′,5′-Cyclic Adenosine Monophosphate
fMLF: N-Formylmethionine-Leucyl-Phenylalanine
FPR1: Formyl Peptide Receptor1
Fsk: Forskolin
IP: Immunoprecipitation
KO: Knockout
MS: Mass Spectrometry
PI: Protease Inhibitor
PLA: Proximity Ligation Assay
PM: Plasma Membrane
PRG: PDZ-RhoGEF
PTX: Pertussis Toxin
TMT: Tandem Mass Tag
WCL: Whole Cell Lysates
WT: Wildtype
YFP: Yellow Fluorescent Protein

## Acknowledgements

We would like to thank U of M Pathology core, especially Venkatesha Barsur, Alexey Nesvizhskii, and Kevin Conlon for running mass spec samples as well as for advice on the data analysis. Srilakshmi Yalavarthi and Gautam Sule for providing isolated neutrophils for this study. We would like to thank Sundeep Malik for BioID2-CaaX cDNA, advice and scientific discussions and John Tesmer for full length PDZ-RhoGEF cDNA mammalian expression construct used in this study.

## Author Contributions

Conceptualized and designed the study: *NRC, AVS.* Participated in research design: *NRC, CAP, AVS.* Conducted experiments: *NRC, SA, SS.* Performed data analysis: *NRC, AVS.* Contributed to the writing of the manuscript: *NRC, AVS.*

## Data Availability

All study data are included in the article and SI Appendix.

