## Supplemental figures, Table and Video legends for "Identification of G Protein α_i_ Signaling Partners by Proximity Labeling Reveals a Network of Interactions that Includes PDZ-RhoGEF"

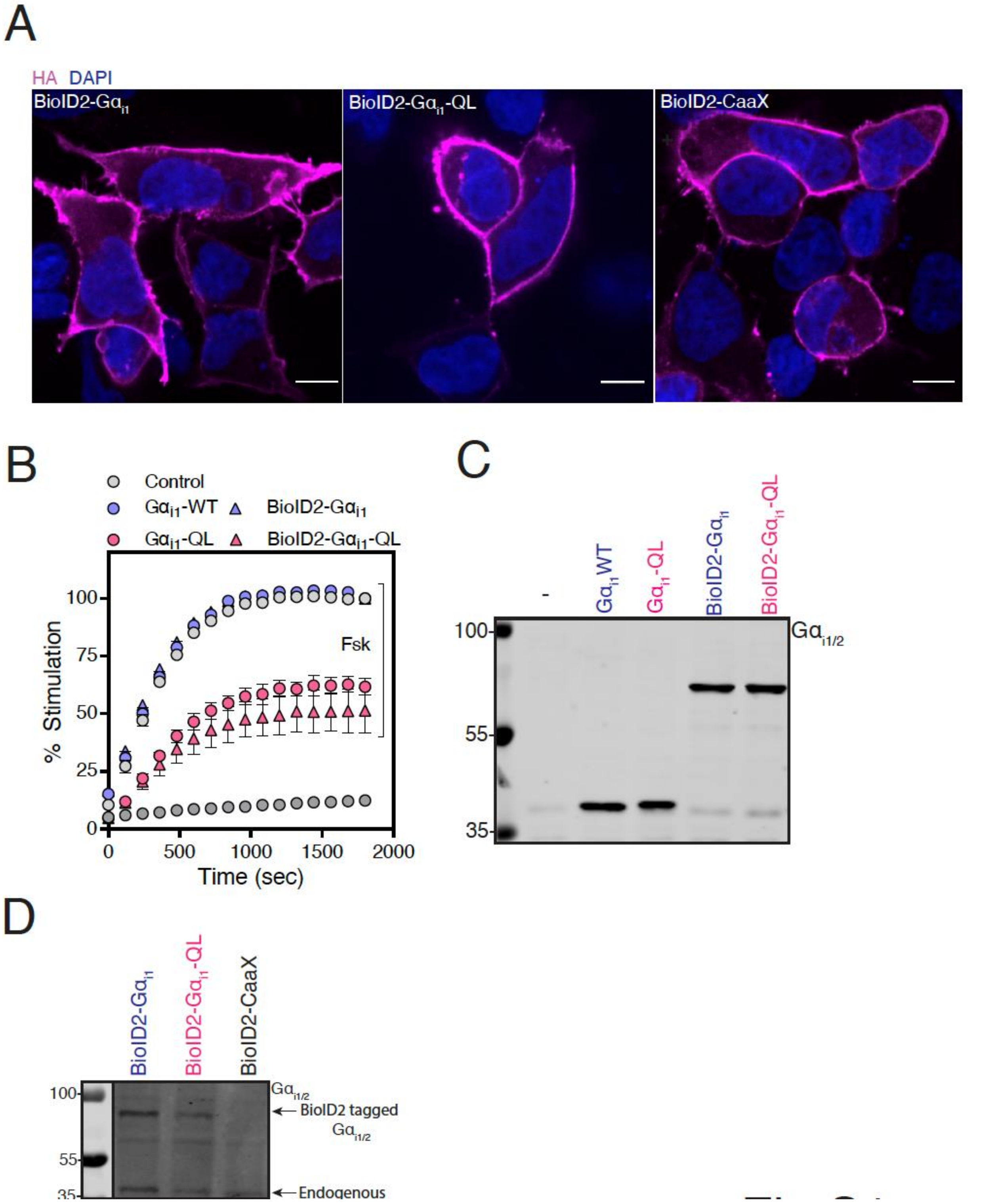


**Supplemental Figure 1. Characterization of BioID2 Fused $G\alpha_{i1}$ and $G\alpha_{i1}$-QL. (A)** BioID2 fused proteins localize predominantly to the PM. Following transfection of HA-tagged BioID2 fused constructs into A293 cells for 48 hr, cells were fixed, permeabilized, and immunostained using an anti-HA antibody. Nuclei were stained with DAPI. Three random fields were imaged, and representative images are shown. Scale bar, 10 µm. **(B)** BioID2 fused $G\alpha_{i1}$-QL is active and inhibits cAMP accumulation. Cells were transfected with cAMP Glosensor$^{TM}$ along with BioID2 tagged and untagged $G\alpha_{i1}$ constructs for 24 hr. Luminescence as a measure of cAMP accumulation was monitored for 30 min (x-axis) after Forskolin (Fsk) stimulation and represented as % stimulation (y-axis) relative to the maximum signal in the respective WT group with 1 µM Fsk treatment. **(C)** Western blot showing relative expression of various constructs. **(D)** BioID2- fused $G\alpha_{i1}$ cDNAs express $G\alpha_{i1}$ protein at a level comparable to endogenous $G\alpha_{i}$ in HT1080 cells used for proximity labeling experiments. Microscopy images, cAMP reporter assay, and western blots represent one of three independent experiments that yielded similar results.

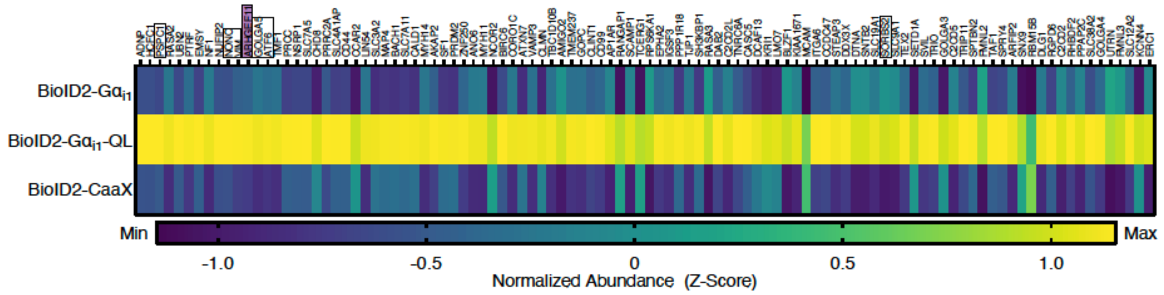


**Supplemental Figure 2. Heat Map of Proteins Identified as Enriched in BioID2-QL Samples Based on the Criteria in Fig. 2C.** Candidate proteins validated in this study are boxed.

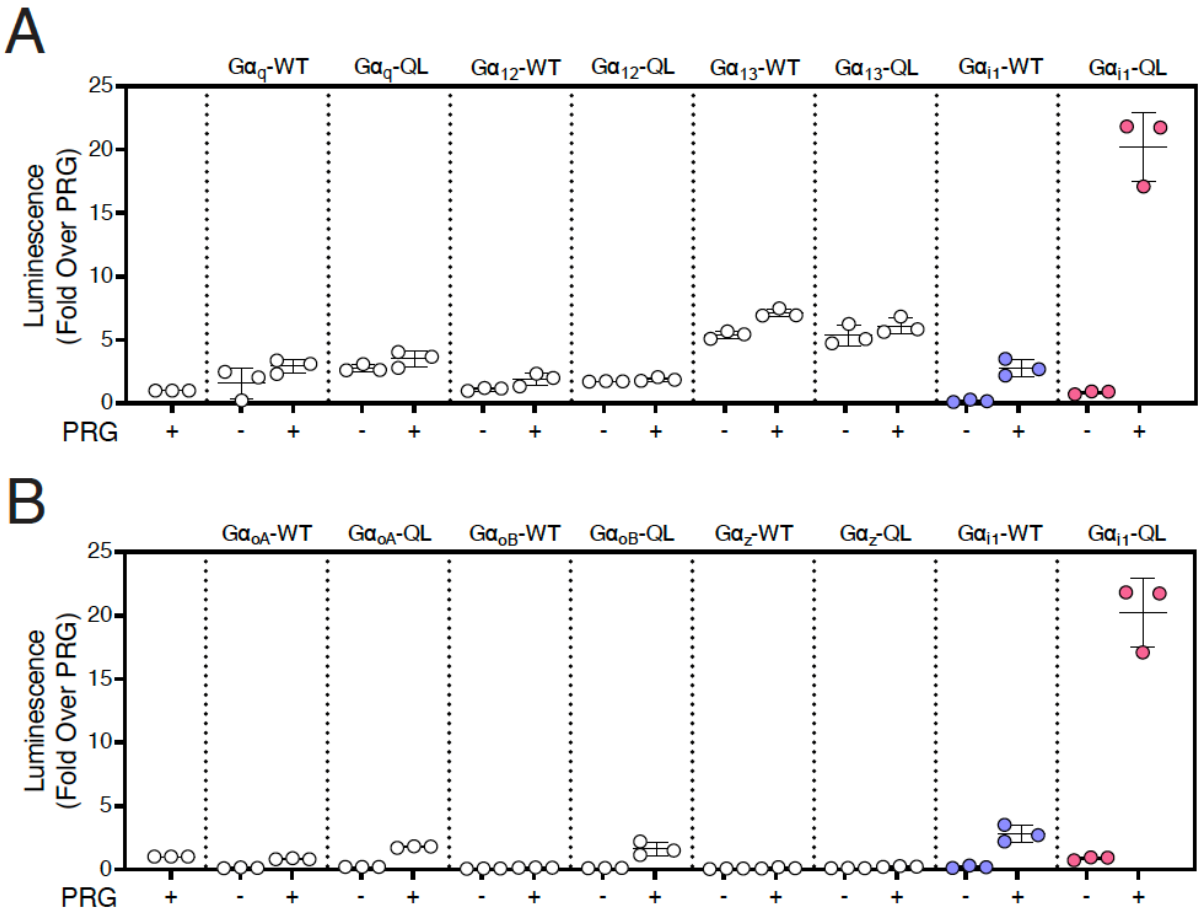


**Supplemental Figure 3. Specificity of PRG Activation by Different G Protein Subunit Family Members in the SRE-luciferase Assay. (A-B)** Each of the indicated Gα protein subunits was transfected with PRG SRE-Luc and assayed as in Fig 4A. The data represent one of three independent experiments all yielding similar results.

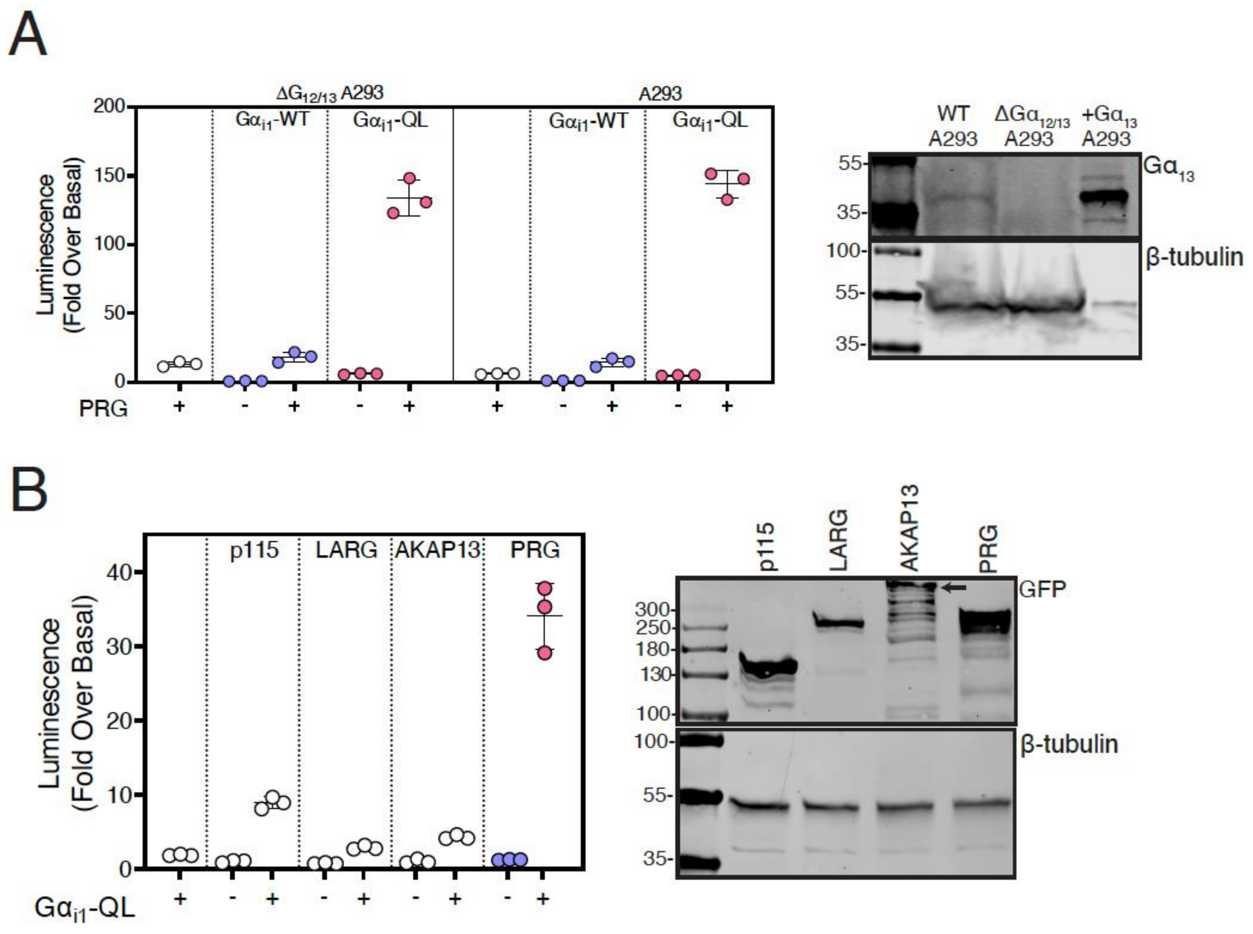


**Supplemental Figure 4. G$\alpha_{i1}$-QL is Specific to PRG Relative to Other RhoGEFs and Does Not Require G$\alpha_{12/13}$. (A)** G$\alpha_{i1}$-QL activates PRG in both A293 and ΔG12/13 A293 cells. Left panel: A293 or $\Delta G_{12/13}$ A293 cells were transfected with SRE-Luc with either G$\alpha_{i1}$ or G$\alpha_{i1}$-QL and assayed as in Fig 4A. Right panel: Western blot showing endogenous G$\alpha_{13}$ expression in both the cell lines and A293 cells transiently expressing G$\alpha_{13}$. β-tubulin was used as an internal control. **(B)** G$\alpha_{i1}$-QL robustly activates PRG and weakly activates other Rho GEFs. Left panel: A293 cells were transfected with SRE-Luc, the indicated RH-RhoGEF family members and either G$\alpha_{i1}$ or G$\alpha_{i1}$-QL and assayed as in Fig 4A. Right panel: Representative western blot showing relative expressions of various GFP tagged RhoGEFs in A293 cells. The data represent one of three independent experiments all yielding similar results.

**Movie legends** for 1) $G\alpha_{i1}$-WT+PRG- DMSO, 2) $G\alpha_{i1}$-WT+PRG- fMLF, 3) $G\alpha_{i1}$-WT+PRG+ PTX-DMSO, 4) $G\alpha_{i1}$-WT +PRG + PTX- fMLF.

Stimulation of A293-FPR1 cells expressing $G\alpha_{i1}$-WT+PRG, with fMLF increased the number of cells with protrusions and PTX treatment strongly inhibited the response. µ-slide 8 well glass bottoms coated with 5 µg/ml fibronectin overnight at 4°C. A293-FPR1 cells were plated on fibronectin and transfected with different constructs, along with YFP, for 48 hours. Cells were imaged at 10x magnification at 20 sec intervals for 40 min. Videos are saved as 10 frames/sec. The cells with protrusions were counted as described in the SI methods.

Movie 1. DMSO treated A293-FPR1 cells expressing $G\alpha_{i1}$-WT+PRG
Movie 2. fMLF treated A293-FPR1 cells expressing $G\alpha_{i1}$-WT+PRG
Movie 3. DMSO treated A293-FPR1 cells expressing $G\alpha_{i1}$-WT+PRG, pretreated with PTX
Movie 4. fMLF treated A293-FPR1 cells expressing $G\alpha_{i1}$-WT+PRG, pretreated with PTX

**Table legends for Table S1, S2, S3**
**Table S1:** Tab 1- GPCRs and Tab 2- All the Receptors Identified by MS experiment detailing their abundance values, ratios, etc. Related to Fig. 2.

**Table S2:** List of 104 proteins identified by MS and selected using the criteria descried in Fig. 2C. Related to Fig. 2 and S2.

**Table S3:** List of Go categories identified for 104 proteins by DAVID software. Related to Fig. 2.
